# Ribosomal Protein Large subunit RPL6 modulates salt tolerance in rice

**DOI:** 10.1101/2020.05.31.126102

**Authors:** Mazahar Moin, Anusree Saha, Achala Bakshi, M. S. Madhav, P B Kirti

## Abstract

The extra-ribosomal functions of ribosomal proteins RPL6 and RPL23a in stress-responsiveness have emanated from our previous studies on activation tagged mutants of rice screened for water-use efficiency (Moin *et al*., 2016a). In the present study, we functionally validated the *RPL6*, a Ribosomal Protein Large subunit member for salt stress tolerance in rice. The overexpression of *RPL6* resulted in tolerance to moderate (150 mM) to high (200 mM) levels of salt (NaCl) in rice. The transgenic rice plants expressing *RPL6* constitutively showed better phenotypic and physiological responses with high quantum efficiency, accumulation of more chlorophyll and proline contents, and an overall increase in seed yield compared with the wild type in salt stress treatments. An iTRAQ-based comparative proteomic analysis revealed the high expression of about 333 proteins among the 4,378 DEPs in a selected overexpression line of *RPL6* treated with 200 mM of NaCl. The functional analysis showed that these highly expressed proteins (HEPs) are involved in photosynthesis, ribosome and chloroplast biogenesis, ion transportation, transcription and translation regulation, phytohormone and secondary metabolite signal transduction. An *in silico* network analysis of HEPs predicted that RPL6 binds with translation-related proteins and helicases, which coordinately affects the activities of a comprehensive signaling network, thereby inducing tolerance and promoting growth and yield in response to salt stress. Our overall findings identified a novel candidate, RPL6 whose characterization contributed to the existing knowledge on the complexity of salt tolerance mechanism in plants.

## 1. INTRODUCTION

Among several abiotic stresses, soil salinity is emerging to be one of the crucial factors having detrimental impact on crop production and it is particularly known to affect photosynthesis by disrupting chloroplast function and stomatal conductance. Moreover, it also affects metabolism, protein synthesis and severe stress can even threaten the very survival of the plants. Salt stress refers to the presence of excess Na^+^ and Cl^-^ ion contents in the soil or the medium in which the plant is growing. The toxicity of salt is considered to be more deleterious than any other agent for a glycophyte like rice as it induces both osmotic stress and ion toxicity (Hu *et al*., 2006, James *et al*., 2011). Because of the accumulation of high sodium ion content during salt stress, water absorption by roots is reduced and the rate of transpiration through stomata is increased resulting in membrane damage, impairment of redox detoxification and overall decrease in photosynthetic activity (Munns, 2005; Rahnama *et al*., 2010). Thus, initial growth suppression of plants occurs due to hyperosmotic effects and subsequent growth arrest is due to the toxic levels of ions.

Salt stress also induces enzyme inhibition and ROS accumulation, which cause DNA damage (Saha *et al*., 2015). The mechanism of salt stress tolerance in plants is complex and has been shown to occur at three levels involving osmotic tolerance (reduction in shoot growth), ion-exclusion (transporter-mediated exclusion of ions by roots) and tissue tolerance (inter and intra cellular ionic compartmentalization) (Tester & Davenport, 2003; Roy *et al*., 2014). Each of these processes occurs either mutually or in isolation and are regulated by a network of candidate genes (Munns *et al*., 2012). Manipulation of the expression of these genes has resulted in transgenic plants with improved tolerance to salt stresses (Park *et al*., 2001; Mukhopadhyay *et al*., 2004; Hu *et al*., 2006; Chen *et al*., 2014; Lee *et al*., 2017; Jiang *et al*., 2019).

Ribosomal proteins are known to play an integral role in generation of rRNA structure and forming protein synthesizing machinery. They are also crucial for growth and development of all organisms (Ishii *et al*., 2006). Since salt stress can result in modification of protein synthesis, it has been observed in many instances that upregulation of genes encoding ribosomal proteins in plants under stressed conditions led to efficient reconstruction of protein synthesizing machinery in cells (Fatehi *et al*., 2012; Omidbakhshfard *et al*., 2012). Ribosomal proteins provide structural stability to the ribosomal complex and participate in protein translation in association with a network of other proteins. Each RP is encoded by a gene that exists as multiple expressed copies in the genome (Degenhardt & Bonham-Smith, 2008).

Several ribosomal protein genes (including *RPS4, 7, 8, 9, 10, 19, 26; RPL2, 5, 18*, and *44*) were among the early responsive genes to be significantly up-regulated during salt stress in salt tolerant Pokkali variety of rice (Dai *et al*., 2005; Sahi *et al*., 2006). Salt stress-dependent up-regulation of plastidial ribosomal protein gene, *PRPL11* has been shown to be involved in improved photosynthetic performance during initial phase of salt stress (Sahi *et al*., 2006). Elevated transcript levels of RPS genes, *S20, S24* and RPL, *L34e* were also observed under NaCl treatment in the root tissues of *Tamarix hispida* (Li *et al*., 2009). Root cDNA libraries of salt stressed corn and rice samples evidenced ribosomal protein transcripts among the abundantly Expressed Sequence Tags (ESTs) (Bohnert *et al*., 2001). Also,a remarkable number of RP genes including *RPL34B, RPL23A, RPS24A, RPS13C, RPS6A, RPL9A* were induced in yeast immediately after the onset of salt stress (Bohnert *et al*., 2001). These analyses point to the fact that the synthesis of ribosomes and an increase in the transcript levels of Ribosomal Protein genes are essential for efficient protein turnover and reconstruction of the protein synthesizing machinery under stress as an immediate consequence of salt shock.

In humans, DNA damage induced by environmental stresses has been shown to recruit RPL6 from the nucleoli to the nucleoplasm at DNA damage sites in a Poly-(ADP-ribose) polymerase-1 and 2-dependent manner where it interacts with the histone protein, H2A/H2AX and promotes its ubiquitination (Yang *et al*., 2019). This H2Ak15ub is necessary to further recruit other repair proteins such as BRCA1 and 53BP1 that promote homologous recombination (HR) and non-homologous end joining repair (NHEJ), respectively (Mattiroli *et al*., 2012; Chapman *et al*., 2013; Fradet-Turcotte *et al*., 2013; Bai *et al*., 2014). In all these signaling cascades, RPL6 has been found to be an important member that is rapidly recruited at DNA damage sites. Depletion of RPL6 results in abrogation of H2A ubiquitination and hence, recruitment of MDC1, BRCA1 and 53BP1 proteins resulting in defects in DNA damage repair (Yang *et al*., 2019). RPL6 also has a role in G2-M checkpoint with decline in its levels resulting in G2-M defects causing damaged cells to enter into mitosis (Yang *et al*., 2019). In addition to RPL6, RPL8 and RPS14 are also shown to be recruited to DNA damage sites (Yang *et al*., 2019).

In rice, the involvement of RP genes in stress-responsiveness has emanated when two of the ribosomal protein encoding genes, *RPL6* and *RPL23A* became activated in the gain-of-function activation tagged mutants of rice screened for enhanced water-use efficiency (Moin *et al*., 2016a). Subsequently, a detailed transcript analysis of all the members of this gene family and functional characterization of a few selected genes revealed their possible involvement not only in improving WUE but also amelioration of abiotic stresses in rice (Moin *et al*., 2016b; Saha *et al*., 2017; Moin *et al*., 2017). A significant up-regulation of considerable number of RPL and RPS genes was noticed both under biotic and abiotic stress conditions. The transcript levels of *RPL6, L7, L18p, L22, L23A, L37* and *RPS6A, S4, S13A*, and *S18A* were found to significantly up-regulated immediately after the application of NaCl treatment (5 min after exposure) and their levels remained significantly high even after prolonged exposure (up to 60 h) (Moin *et al*., 2016a).

In the present study, we report on the detailed characterization of *RPL6* by its constitutive expression in *indica* rice followed by studies on phenotypic and physiological responses of overexpressing lines under varied levels of salt (NaCl) stress. Further, we present a complete protein profile of an high expression line of *RPL6* treated with NaCl using a quantitative proteomic approach (iTRAQ). We have identified proteins that were highly expressed in *RPL6* treated with NaCl and also discussed their possible functional association in improving salt stress tolerance. Our collective results show that *RPL6* balances the growth and yield under salt stress by highly expressing proteins associated with growth, development and stress signaling pathways, thereby inducing salt tolerance.

## 2. MATERIALS & METHODS

### 2.1. Design of *RPL6* binary vector

After the identification of the involvement of *RPL6* in enhancing water-use efficiency (WUE) in rice (Moin *et al*., 2016a), the full-length cDNA sequence (681 bp) of the *RPL6* gene (LOC_Os04g39700) was obtained from the Rice Genome Annotation Project database (RGAP-DB). The retrieved sequence was further verified through nucleotide and protein BLAST searches in RAP-DB, NCBI and Hidden Markov Model (HMM) of Pfam databases. When the sequence of *RPL6* from all the databases showed identical matches, the primers were synthesized by incorporating *Xho*I and *Nco*I restriction sites at the forward and reverse ends of the cDNA sequence, respectively for subsequent cloning steps. The *RPL6* sequence was PCR amplified from the cDNA of BPT-5204 rice variety. Initially, the pRT100 (Addgene, A05521) vector was double digested with *Xho*I and *Nco*I restriction enzymes. In the next step, the *Xho*I and *Nco*I-treated *RPL6* sequence was ligated into the double-digested pRT100 vector. This step brought the *RPL6* in a transcriptional fusion with CaMV35S promoter and poly-A tail at its 5’ and 3’ ends, respectively resulting in the *RPL6* plant expression cassette. The entire expression cassette with the promoter and poly-A tail was removed as a *Pst*I fragment and cloned into the binary vector, pCAMBIA1300. This binary vector (pCAMBIA1300: *RPL6*) carrying expression cassette of *RPL6* was mobilized into the *Agrobacterium tumefaciens* strain EHA105 for genetic transformatioof rice.

### 2.2. *In planta* transformation of 35S: *L6* in *indica* rice

The BPT-5204 (Samba Mahsuri), a very widely cultivated *indica* rice variety has been used to develop transgenic plants overexpressing *RPL6*. The pCAMBIA1300: *RPL6* (which is referred to as 35S: *L6* hereafter in this manuscript) binary vector was transformed into rice using *in planta* method of transformation as reported previously (Moin *et al*., 2016a). In short, after surface sterilization of BPT-5204 seeds with 4% sodium hypochlorite (20 min) followed by five washes with sterile double-distilled water, they were soaked in water overnight (12-16 h) to allow the embryonic elongation to occur. Following this, a sterile needle that was dipped in *Agrobacterium* suspension containing 35S: *L6* construct was gently pierced at the base of the embryo which would later produce hypocotyl and subsequently cotyledons. The infected seeds were subjected to a vacuum of 15 mmHg for 20 min, after which the vacuum was released suddenly. This process of vacuum infiltration coupled with the sudden release of vacuum forces the *Agrobacterial* cells to replace the intercellular air spaces of the explant. Appropriate infection stage of the seeds and vacuum infiltration are the crucial factors determining the efficiency of *in planta* transformation in rice. The transformation efficiency of the 35S: *L6* transgenic plants was nearly 20%. After initial germination, the *Agrobacterium* infected seeds (which were considered as T_0_) were transferred to black alluvial soil in the pots maintained under controlled greenhouse conditions (30 ± 2°C with 16/8 h of light/ dark photoperiod).

### 2.3. PCR, Southern-blot hybridization and quantitative real-time PCR (qRT-PCR)

The plants obtained from each of *Agrobacterium*-infected seed were considered as a separate line owing to the independent integration of the T-DNA. After harvesting T_0_ plants, T_1_ seeds were collected separately and allowed to germinate on MS selection medium containing the antibiotic, Hygromycin (50 mgl^−1^). The 35: *L6* plasmid worked as a Positive Control (PC), whereas the plants that were obtained from the non-germinated seeds on the selection medium followed by rescue on the selection-free medium were used as Negative Control (NC). The T-DNA of the 35S: *L6* binary vector contains the expression cassettes of *L6* and *hpt*II, which acts as a plant selection marker. Because *L6* is endogenous to rice, *hpt*II was used to confirm the transformed plants in subsequent generations. To confirm the transgenic nature of transformed plants and also to identify the T-DNA copy number, the transformed plants were subjected to Southern-blot hybridization as per the standard protocols.

Total RNA was isolated using standard Trizol method (Sigma-Aldrich, US) from the root and shoot tissues of two-week-old seedlings of NC and 35S: *L6* transgenic lines before and after treatment with different concentrations of NaCl. About 2 μg of the isolated RNA was used to synthesize the first-strand cDNA using reverse transcriptase enzyme (Takara Bio, Clontech, USA). The cDNA was diluted seven times with sterile Milli-Q water (1:7 ratio). After gentle pipetting, 2 μl of the diluted cDNA was used in qRT-PCR experiments to analyze the transcript level of various genes.

### 2.4. Evaluation of seedlings for salt tolerance

Based on the results of semi-Q and qRT-PCR, T_3_ seedlings from three low and three high expression lines were subjected to salt stress screening along with NC. The seeds obtained from NC and each of these six lines were germinated on solid MS medium for two weeks. The healthy seedlings in replicates of five from NC and each of six transgenic lines were transferred to liquid MS medium containing three different concentrations of NaCl solution such as 100 mM (low), 150 mM (medium) and 200 mM (high) at pH 5.8. Salt treatment at each concentration was applied for a duration of three and five days following which all the seedlings along with NC were allowed to recover by transferring to salt-free (solid MS without NaCl) medium.

### 2.5. Transcript analysis of *L6*

To check the transcript levels of the *L6* gene in T_3_ transgenic lines, semi-Q and qRT-PCR were performed. Semi-Q PCR was conducted with an initial denaturation at 94 °C for 1 min followed by 28 repeated cycles of 94 °C for 30 s, *L6* annealing temperature of 56 °C for 25 s and an extension temperature of 72 °C for 45 s. The PCR reaction was terminated with a final extension step at 72 °C for 5 min. Rice specific actin, *act1* was used as an internal reference gene. The transcript levels of the *L6* gene in transgenic lines were determined based on the intensity of semi-Q PCR bands electrophoresed on 1.5% agarose gel with respect to NC. The transgenic lines with pale bands were considered as low expression lines, whereas those with intense bands were treated as high expression lines. The levels of *L6* transcripts observed in semi-Q PCR were further verified through qRT-PCR using SYBR master mix (Takara Bio, USA). Rice *act1* was used as a house-keeping gene and the cDNA synthesized from NC was used to normalize the expression pattern by ΔΔC_T_ method (Livak and Schmittgen, 2001). The reaction and cyclic conditions for qRT-PCR were the same as used in semi-Q PCR.

### 2.6. Phenotypic studies

The fresh weight and, root and shoot lengths (measured using a 15 cm scale bar) of two-week-old NC and transgenic seedlings were measured before and after treatment with 150 and 200 mM NaCl. In addition, various phenotypic parameters were also recorded after transfer to greenhouse such as total plant height, culm length, tiller length, number of tillers per plant, leaf area, panicle length, number of panicles per plant, number of grains per panicle and the total number of seeds per plant (seed yield). These were measured in NC and six transgenic lines (T_3_ generation) that were grown in the absence of salt (untreated) and revived after 5 d of exposure to 150 and 200 mM salt stress. Each reading was taken at the corresponding growth stage of the plant as a mean of three biological replicates which were plotted as bar diagrams.

### 2.7. Chlorophyll and Proline contents

Proline is an amino acid that acts as an osmolyte and compatible solute in cells. When plants are exposed to environmental stresses, they tend to accumulate proline to a certain level where it exhibits diverse functions such as a metal chelator, protein compatible hydrotrope, ROS detoxification, stabilizing cellular membranes, maintaining protein integrity and appropriate NADP^+^/NADPH ratios (Hare & Cress, 1997; Strizhov *et al*., 1997; Ashraf & Foolad, 2007). These activities of proline after accumulation in response to stresses have been correlated with stress tolerance (Petrusa & Winicov, 1997). To check the proline content and its correlation with the level of salt tolerance in transgenic plants, proline was extracted from leaves of NC and transgenic lines (T_3_ generation) with aqueous sulphosalicylic acid. The isolated proline was treated with ninhydrin and measured at 520 nm (Bates *et al*., 1973).

Exposure of plants to salt stress generates reactive oxygen species that degrade the chlorophyll contents (Verma & Mishra, 2005). This chlorophyll degradation is used as an indicator to assess the extent of oxidative damage that occurred in the cell due to the ongoing stress (Rio *et al*., 2005). The chlorophyll contents (Chl-a and b) were estimated using 100 mg of leaves obtained from NC and transgenic lines ground in 80% acetone. The absorbance of the extracts was measured at 663 and 645 nm and the concentration of pigments were calculated as per standard protocols (Arnon, 1949; Zhang *et al*.,, 2009).

### 2.8. Chlorophyll fluorescence

Chlorophyll Fluorescence (CF) is a physiological parameter that measures the activity of photosystem-II (PSII), also widely used to assess the response of a plant to biotic and abiotic stresses (Murchie & Lawson, 2013). CF, which is a measure of re-emitted light from PSII, gives information related to quantum efficiency and overall photosynthesis and ultimate productivity of a plant. CF is measured empirically as F_v_/F_m_ (where Fv is the variability in fluorescence and Fm is the maximum possible yield of fluorescence resulting from a saturating pulse of 8000 μmol m^−2^ s^−1^). For plants that are healthy and grown under unstressed environments, Fv/Fm is as high as ~0.83, which corresponds to maximum quantum yield. Exposure to stress leads to the inactivation of PSII resulting in a significant reduction of Fv/Fm (Maxwell & Johnson, 2000). In the present study, CF was measured using a portable Pulse Amplitude Modulation (MINI-PAM) instrument (Murchie & Lawson 2013) as per the manufacturer’s protocol (Walz, Effeltrich, Germany). Measurements were recorded in the dark-adapted leaves of NC and six lines of 35S: *L6* plants (T_3_ generation) after four weeks of revival from the application of 200 mM salt stress for five days. Readings were taken in biological triplicates and the mean of Fv/Fm was plotted as a bar diagram.

### 2.9. iTRAQ-based protein identification

Two-week-old rice seedlings of WT and one high expression line of *L6* transgenic rice (*L6-5*) were subjected to 200 mM NaCl treatment for a period of five days. The quantitative proteomic analysis was performed in four seedling samples that include WT-untreated, WT-treated, L6-untreated and *L6*-NaCl using the iTRAQ (isobaric Tags for Relative and Absolute Quantitation) method, which has high degree of sensitivity and provides comprehensive information about the expressed proteins compared to the other techniques. The iTRAQ technique was commercially performed from the proteomic services of Sandor Lifesciences Pvt. Ltd.

### 2.10. Transcript analysis of selected genes

To validate the expression pattern of proteins obtained in iTRAQ analysis on a selected line, transcript levels of some of the highly expressed genes like *OsTOR, OsWRKY51, OsRPL7, OsRPL37* and *OsRPS20* were studied through qRT-PCR. Their transcript levels were checked in *L6* transgenic lines before and after 5 d of exposure to 200 mM NaCl. Before treatment, the transcript levels of these genes in transgenic lines were normalized with untreated NC. Since the NC plants did not recover after treatment with 200 mM NaCl, these transcripts in high expression lines were normalized with respect to a recovered low expression line.

### 2.11. Statistical analysis

All the qRT-PCR experiments were conducted in biological triplicates, whereas the phenotypic characters were sampled in replicates of five plants (NC and transgenic). All the experiments were conducted in six transgenic lines (three high and three low expression lines) belonging to T_3_ generation with respect to their corresponding NC. The SigmaPlot v.11 was used for statistical analysis. Statistical significance was calculated using one-way ANOVA and significance at *P* < 0.05 was represented with asterisks in bar diagrams. The protein networking was constucted using STRING v11, which generates networks based on functional association (Szklarczyk *et al*., 2015). A threshold score of 0.4 which is widely used was also applied in this study. This analysis also provides a score between 0.15 to 0.9 representing the degree of connection between two protein nodes.

## 3. RESULTS

### 3.1. Selection of 35S: *L6* plants

The 35: *L6* vector (Fig. 1a) was confirmed by PCR amplification of *hpt*II and *L6* genes, which produced expected fragment sizes of 1025 and 680 bp, respectively. The vector was also digested with *Pst*I restriction enzyme to release the 1300 bp expression cassette of *L6* corresponding to the 35S promoter (350 bp), *L6* cDNA (681 bp) and poly-A tail (250 bp). After confirmation of 35S: *L6* binary vector and the EHA105 strain carrying the vector, it was transformed into rice through the *in planta* method of genetic transformation. The seeds obtained from the primary infected plants were considered as T_1_, which were advanced to T_2_ and T_3_ generations by allowing them to germinate on MS medium containing 50 mgl^−1^ Hygromycin as the plant selection antibiotic. As the transformed seedlings started to germinate within 3-4 days with subsequent normal healthy growth, they were transferred to pots in the greenhouse and further confirmed with PCR analysis. The non-transformed seedlings became bleached after initial germination for 1-2 days. Some of them were revived by transferring to Hygromycin-free medium and were used as non-transformed control (NC) for a comparative study with transgenic plants. PCR was performed with the *hpt*II gene, which produced a band of 1025 bp in the transformed plants (Fig. 1b). About 500 BPT-5204 seeds were infected with the EHA-105 carrying 35S: *L6*, of which nearly 100 plants (20%) appeared to be positive with Hygromycin selection and PCR amplification. Among the eleven Hygromycin-positive transformants, six were found to be positive through Southern-blot hybridization with a single copy integration of T-DNA (Fig. 1c).

**Figure 1.**
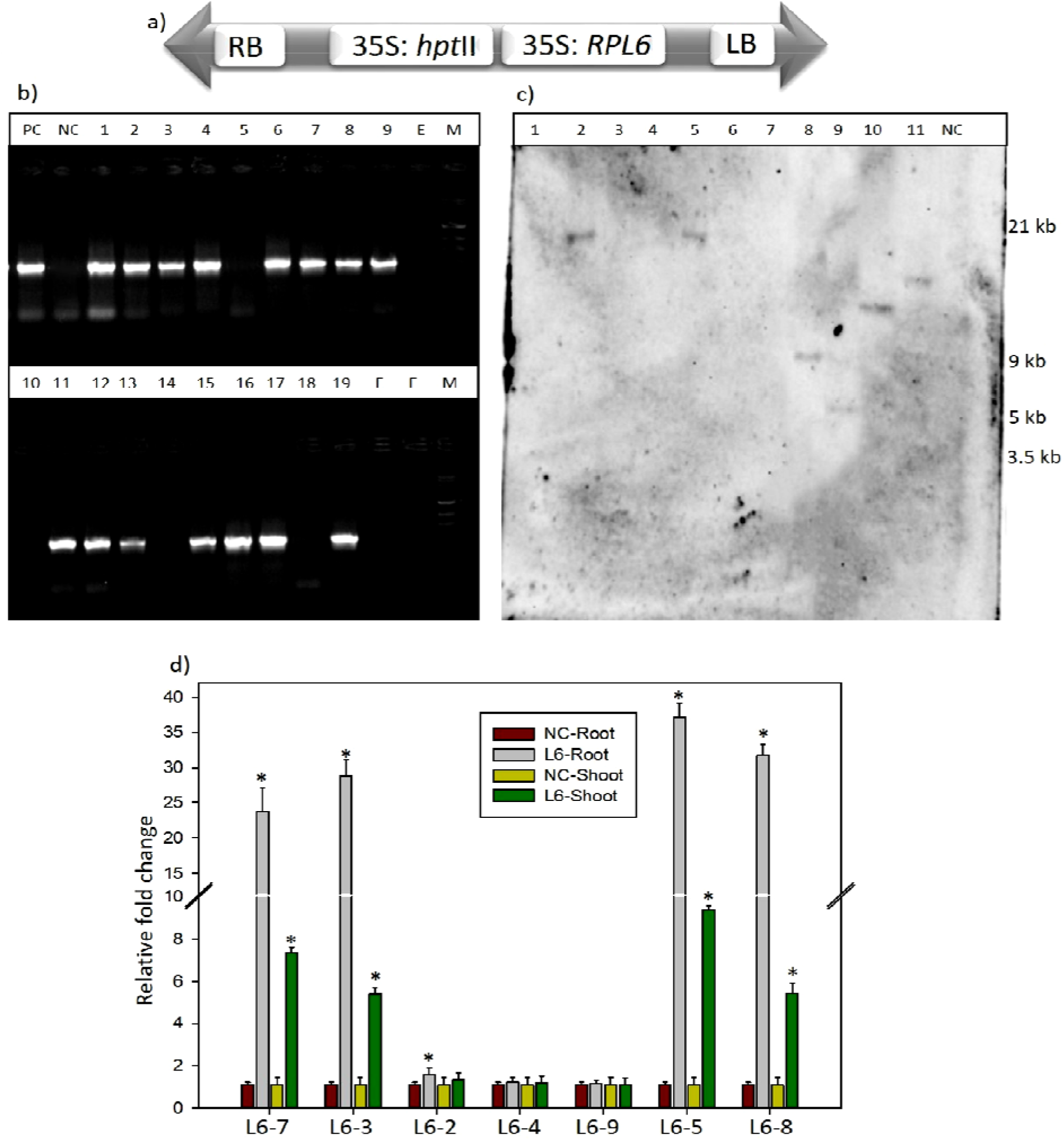
Cloning and molecular investigations of *RPL6*. The entire expression cassette of *RPL6* along with the promoter and poly-A tail as a *Pst1* fragment was cloned into (a) T-DNA of binary vector, pCAMBIA1300. RB and LB, Right and Left borders of the T-DNA, respectively; 35S, CaMV35S promoter; E, empty well; M, λ*EcoR*I-*Hind*III marker. The transformed plants were analyzed using (b) PCR and (c) Southern-blot hybridization. Lanes 1-19 (in PCR) and 1-11 (in Southern-blot) refers to transgenic samples. NC, Negative Control; PC, Positive Control. The sizes labeled in Southern-blot are according to λ *EcoR*I-*Hind*III marker. (d) The *RPL6* transcript levels in transgenic plants were studied using qRT-PCR. Based on the transcript patterns, *L6-3, L6-5, L6-7* and *L6-8* were considered as high expression, and *L6-2, L6-4* and *L6-9* as low expression lines.

### 3.2. Separation of 35S: *L6* lines with semi-Q and qRT-PCR

All the PCR-positive T_3_ seedlings were subjected to semi-Q and qRT-PCR to check the transcript levels of the *L6* gene. Based on the band intensity, four were identified as high expression lines and remaining lines had pale bands (Supplementary Fig. 1). The expression of *L6* in qRT-PCR was studied in root and shoot tissues separately, of which high transcript levels were noticed particularly in roots in all the lines. Because of this, the transcript levels of *L6* in roots were considered for the separation of the lines. The transgenic plants with <2-fold of *L6* transcripts were considered as low-expression lines and the plants with >20-fold were considered as high-expression lines. Based on the qRT-PCR results, three lines, *L6-2, L6-4*, and *L6-9* were identified as low expression lines with transcript levels of 1.5, 1.2 and 1.1 fold in roots. Four lines, *L6-3, L6-5, L6-7* and *L6-8* showed an elevation of gene transcripts >20-fold in roots, while, it was <10-fold in shoots and these were considered as high expression lines. Among all these, two lines, *L6-5* and *L6-8* exhibited the highest transcript level of 37 and ~32-fold in roots with 9 and 5-fold in shoot tissues, respectively (Fig. 1d). Hence, these two lines (*L6-5* and *L6-8*) along with *L6-3* (28-fold) and three low expression lines (*L6-2, L6-4*, and *L6-9*) were selected for a detailed phenotypic and physiological investigations related to salt tolerance.

### 3.3. Salt tolerance in *L6* transgenic seedlings

Seedlings from NC, three high (*L6-3, L6-5* and *L6-8*) and three low expression lines (*L6-2, L6-4*, and *L6-9*) were exposed to low (100 mM), moderate (150 mM) and high (200 mM) concentrations of NaCl in liquid MS medium (devoid of organics) for 3 and 5 d continuously. During exposure to 100 mM NaCl for 3 and 5 d, all the low and high expression lines exhibited normal growth and showed no signs of wilting (Fig. 2a), and the NC started to curl only after 3 d of exposure. All these seedlings along with NC were recovered when shifted to salt-free medium. In the presence of 150 mM NaCl, the tip of NC seedlings started to wilt after 24 h of exposure (Fig. 2b). The seedlings of *L6-2, L6-4* and *L6*-*9* (low expression lines) showed mild signs of wilting after 5^th^ d of exposure. In the case of *L6-3, L6-5* and *L6-8*, no signs of leaf curling were noticed even after 5 d of exposure. When transferred to NaCl-free medium, all the triplicate seedlings of NC, *L6-4* and *L6-9* failed to recover, whereas only one seedling of *L6-2* recovered (Fig. 2c). The lines, *L6-3, L6-5* and *L6-8* recovered completely and continued to grow normally. When exposed to 200 mM for 3 d, NC showed immediate leaf yellowing and curling but other seedlings remained green (Fig. 2d). After 5 d, *L6-3, L6-5* and *L6-8* appeared healthier than low expression lines (Fig. 2e). After transfer to NaCl-free medium, only one low expression line, *L6-2* recovered, whereas all the high expression lines, *L6-3, L6-5* and *L6*-*8* recovered back to normal growth after the removal of the stress solution. The seedlings of NC, *L6-4* and *L6-9* became completely wilted and have not been able to recover.

**Figure 2.**
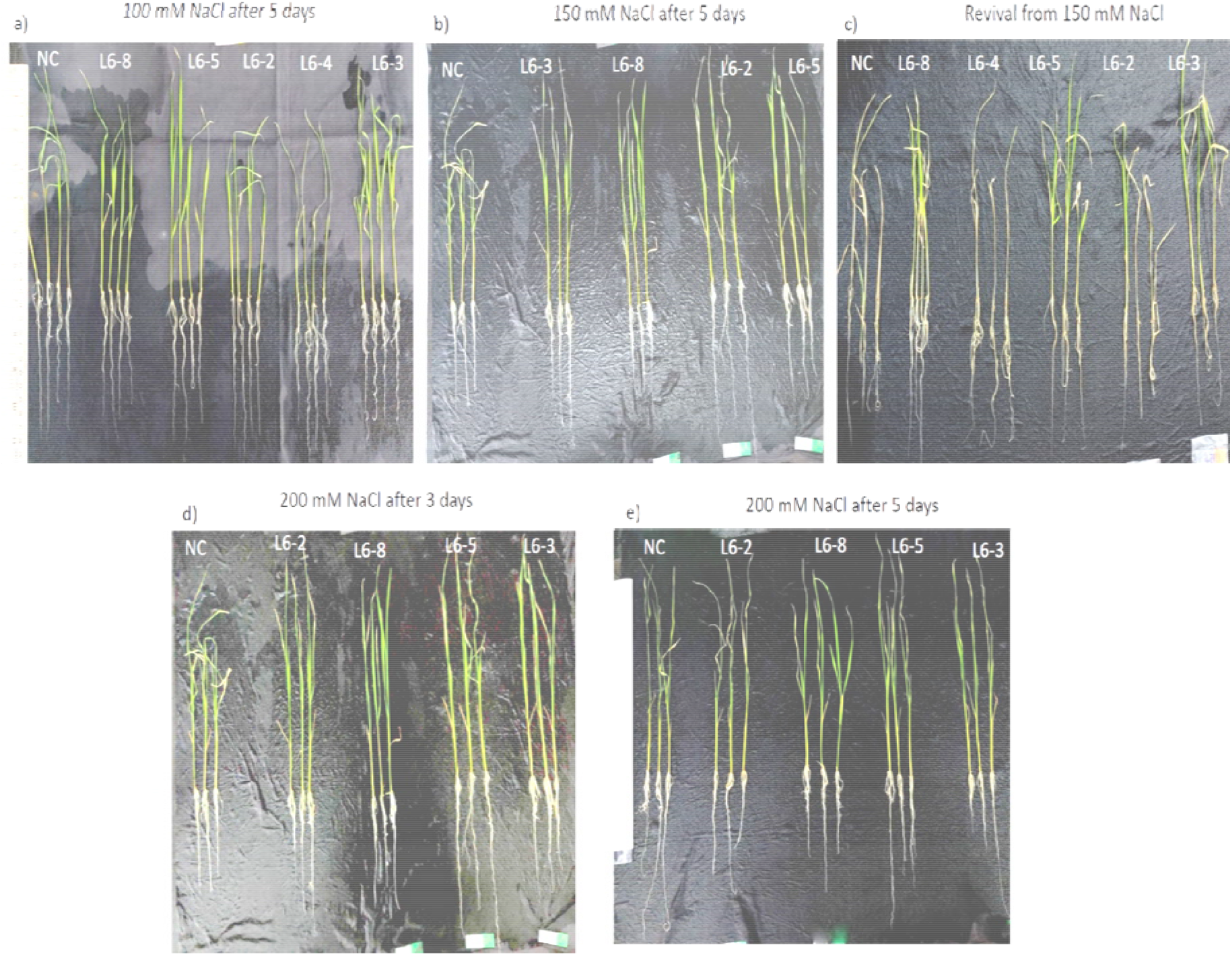
Salt stress analysis of *RPL6* transgenic seedlings. Two-week old *RPL6* transgneic and negative control seedlings were treated with (a) 100 mM, (b, c) 150 mM and (d, e) 200 mM NaCl. Their fresh weight, root and shoot lengths were measured before and after treatments.

### 3.4. Phenotypic analysis

The fresh weight, root and shoot lengths were measured in two-week-old seedlings of NC and six transgenic lines (three high and three low expression lines) before and after 3 d and 5 d of exposure to 150 and 200 mM NaCl. Before transfer to salt stress solution, the mean shoot length of two-week-old NC, low and high expression lines were 16, 15 and 16 cm, respectively (Supplementary Fig. 2a). The mean root lengths of these lines were 7, 6 and 7 cm, respectively, whereas the mean fresh weights were 0.25, 0.21 and 0.25 g, respectively (Supplementary Fig. 2b). After 3 d and 5 d of treatment at 150 mM, the fresh weights ranged from 0.21 (*L6-4*) to 0.24 g (*L6-3*) and 0.24 (*L6-4*) to 0.27 g (*L6-5*), respectively (Supplementary Fig. 2c). After treatment with 200 mM for 3 d and 5 d, the fresh weight of *L6-5* was highest with 0.26 and 0.29 g, respectively. After 5 d, all these seedlings were shifted to NaCl-free medium. Among six transgenic lines, *L6-4* and *L6-9* failed to recover 5 d after 150 mM and 3 d after 200 mM treatment to NaCl. After one week of recovery in NaCl-free MS medium, all the recovered seedlings were shifted to pots in the greenhouse for further phenotypic and physiological characterization. Because NC did not recover after salt stress, the analyses of transgenic plants hereafter were made with respect to the wild type (WT) BPT-5204 rice.

Under normal (NaCl-free) conditions, the yield-related parameters such as the size and number of tillers and panicles, and total seed yield of high expression lines were similar to WT. The total seed yield in both the untreated WT and three high expression lines was ~26 g (185 seeds). In transgenic lines recovered from 5 d of exposure to 150 mM NaCl, the number of tillers ranged from 3 (*L6-2*) to 5 (*L6-5*) per plant, the panicle size was between 10 (*L6-2*) to 15 cm (*L6-5*), number of grains per panicle was between 30 (*L6-2*) to 45 (*L6-5*) and the total seed yield was 12 (*L6-2*) to 21 g (*L6-5*) per plant. After recovery from 150 mM, the seed yield in three high expression lines, *L6-3, L6-5* and *L6-8* was 16, 21 and 19 g per plant, respectively. The total yield in *L6-5* after 5 d of exposure to 150 mM NaCl was a little less than WT rice grown under salt-free conditions (untreated) whose yield was 26 g. After 200 mM treatment, the number of tillers per plant ranged from 1 (*L6-2*) to 4 (*L6-5*), panicle size was between 6 (*L6-2*) to 13 cm (*L6-5*), the number of panicles per plant were 2 (*L6-2*) to 3.5 (*L6-5*) and the number of grains per panicle was 14 (*L6-2*) to 42 (*L6-5*). The total seed yield was between 3 to 16 g in a low (*L6-2*) and high (*L6-5*) expression lines, respectively. After recovery from 5 d of exposure to 150 and 200 mM NaCl, the total seed yield in the high expression line, *L6-5* was 21 and 16 g, respectively (Supplementary Fig. 3). Based on these observations, *L6-5* was found to be a high performing line even after exposure to high concentrations of salt. These results are suggestive of the fact that 35S: *L6* has an important role in inducing salt tolerance in rice with a little compromise on the total yield of the crop.

### 3.5. Accumulation of chlorophyll and proline contents

The contents of total chlorophyll, Chl-a, and b were measured in WT, low expression and three high expression lines. Under normal conditions (NaCl-free), the levels of Chl-a and total chlorophyll were >10 μg/ml in all these plants. The concentration of Chl-b in WT and high expression lines remained the same (~5 μg/ml), but it was slightly reduced in low expression lines (3 μg/ml). After recovery from 200 mM salt stress, all the three chlorophyll contents were significantly elevated in *L6-5*, moderately elevated in *L6-3* and *L6-8* and decreased in *L6-2* line (Fig. 3a-c). The high chlorophyll contents are associated with high quantum efficiency and hence, an increased photosynthetic activity played an imported role in sustainable yield in high expression lines. Under normal conditions, the proline content in NC, high expression (*L6-5*) and a low expression line (*L6-9*) was 0.5, 0.9 and 0.3 μg/mg, respectively. After treatment with NaCl, the proline content in transgenic lines was increased to more than 1-fold, indicating that cytosolic osmotic potential is preserved under salt stress in transgenic lines (Fig. 3d).

**Figure 3.**
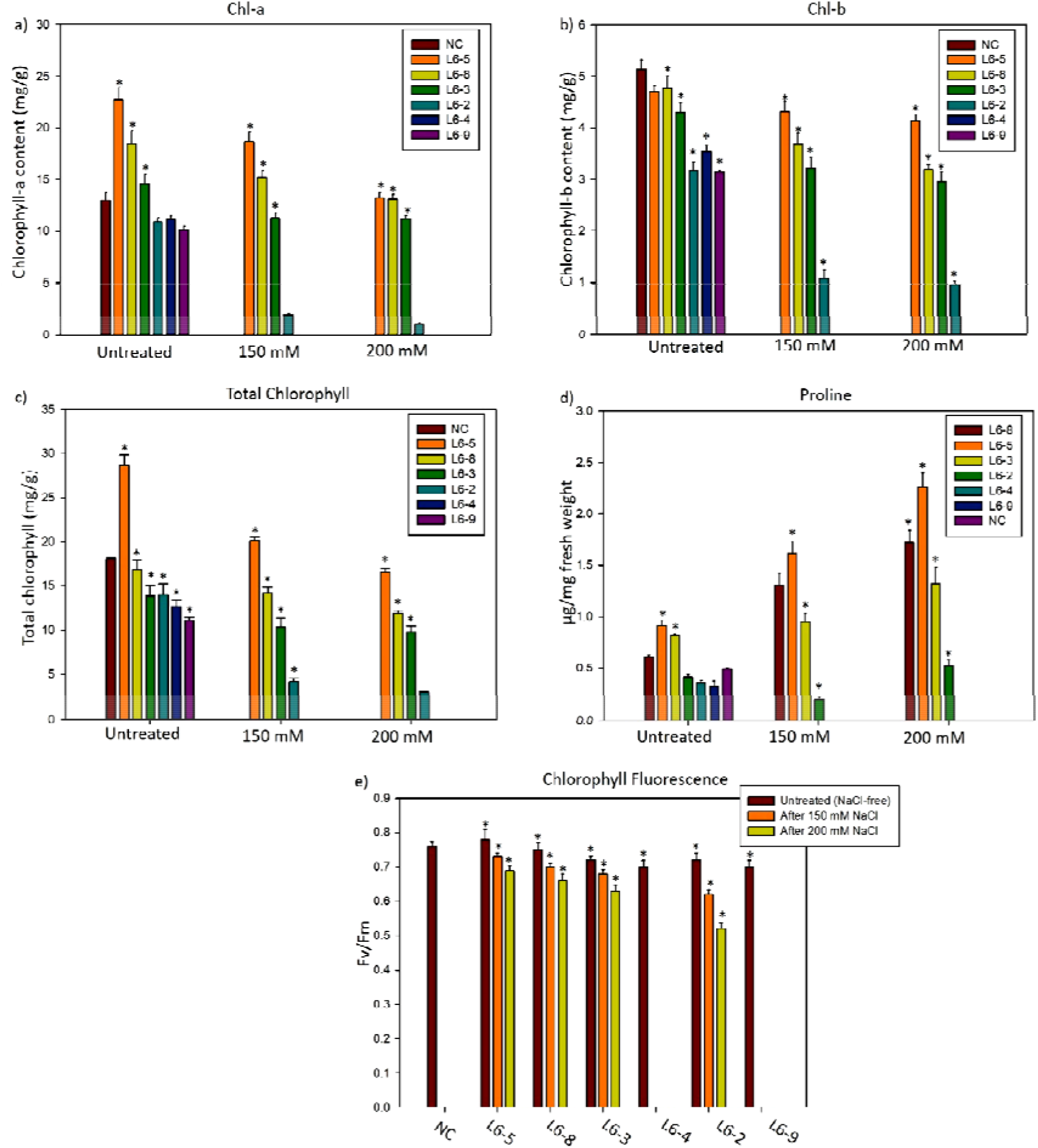
Physiological analysis of *RPL6* transgenic plants. The contents of (a) chlorophyll-a, (b) chl-b, (c) total chlorophyll and (d) proline were measured in six *RPL* transgenic and negative control plants before and after treatment with 150 and 200 mM NaCl. The (e) chlorophyll fluorescence was performed with MINI-PAM. Mean values of physiolgical data with ± standard error represented with asterisks were considered statistically significant at *P* < 0.05.

### 3.6. Quantum efficiency of PSII

The quantum efficiency of PSII was measured using a portable MINI-PAM in dark-adapted leaves of transgenic lines that were grown under normal conditions without applying salt stress (untreated) and those that were recovered after the application of 150 and 200 mM NaCl. Under untreated conditions, the CF of WT rice was 0.76, whereas the CF of transgenic lines ranged from 0.70 (*L6-9*) to as high as 0.78 (*L6-5*). After the application of 150 and 200 mM NaCl, the quantum efficiency in a solely recovered low expression line, *L6-2*, was 0.62 and 0.52, respectively. After 5 d of NaCl treatment at 150 mM, the CF in three high expression lines, *L6-3, L6-5* and *L6-8* was 0.68, 0.73 and 0.70, respectively, while the quantum efficiency in these three lines after 5 d of exposure to 200 mM was 0.63, 0.69 and 0.66, respectively (Fig. 3e).

### 3.7. iTRAQ proteome analysis

The iTRAQ quantitative proteome approach was employed in WT-untreated, WT-NaCl, *L6*-untreated and *L6*-NaCl seedlings. In total, 4,378 differentially expressed proteins (DEPs) were identified in all the four samples. About 35% (1,507) of these exhibited a coverage of >10%, indicating high confidence. The molecular weight of 65% of DEPs was in the range of 10-100 KDa and 35% had >100 KDa, again indicating a good coverage. Of the total DEPs 2,856 (66%) proteins were up-regulated with >1-fold and 1, 522 (34%) were down-regulated (<1-fold) in *L6*-NaCl treated line. Further, among the up-regulated ones, we have identified about 333 proteins in *L6*-NaCl line whose expression was >1.3-fold and higher than WT-untreated, WT-NaCl and *L6*-untreated samples and these were considered as Highly Expressed Proteins (HEPs). The high expression of these proteins might have been controlled by RPL6 in response to NaCl stress, but not by NaCl stress alone as they were highly expressed particularly in *L6*-NaCl line and not in other three samples. The HEPs were categorized into different groups based on their biological functionas such as proteins with catalytic activities, transporters, transcription factors, heat shock proteins (HSPs) and translation-related proteins. The catalytic proteins were further categorized into transferases, oxidoreductases, lyases, ligases, hydrolases, peptidases, dehydrogenases, kinases, demethylases, phosphatases, DNA-dependant and RNA-dependant catalytic proteins. Catalases accounts for 64% of the total HEPs, followed by transporters (20%), transcription factors (10%), translation-related proteins (2%) and HSPs (1%). We have provided an emphasis on these 333 proteins with respect to their role in development, yield and stress responses and also predicted their network with RPL6 *in silico*. The functional grouping of the total DEPs and HEPs were also provided in the form of pie-charts in Supplementary Figs. 4 & 5, respectively. The detailed list of the HEPs were provided in Supplementary Table. 1.

#### 3.7.1. Catalytic activities

##### 3.7.1.1. Transferases

Transferases catalyze the transfer of functional groups from one molecule to a recipient. About 36 diverse transferases constituting 17% of the total catalases, which are involved in carbohydrate, DNA and amino acid-mediated metabolic processes were highly expressed with >1.3 FC in the *L6*-NaCl line indicating the involvement of this protein in maintaining the cellular metabolism under salt stress conditions (Fig. 4a). Some of the transferases with the highest expression include Hydroxy-cinnamoyl transferase 1 followed by UDP-glycosyl transferase 79, glucosamine 6-phosphate N-acetyl transferase 2, Acyl transferase 1, diacylglycerol O-acyltransferase 1-1, indole-3-acetate O-methyl transferase 1, glucose-1-phosphate adenylyl transferase small subunit 2, cycloartenol-C-24-methyltransferase 1, glucose-1-phosphate adenylyl transferase large subunit 1.

**Figure 4.**
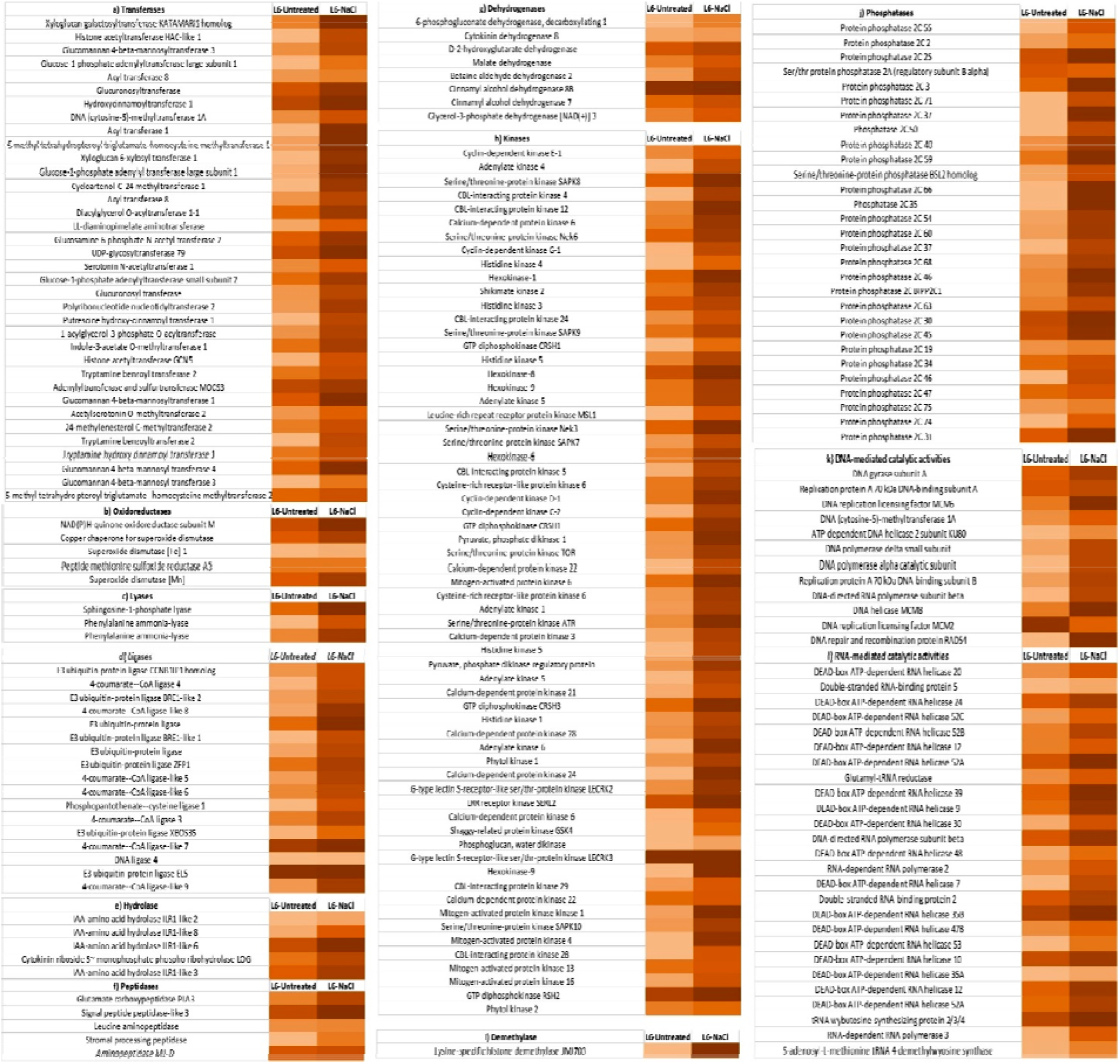
Expression pattern of catalases. The level of expression of different catalases like (a) transferases, (b) oxidoreductases, (c) lyases, (d) ligases, (e) hydrolases, (f) peptidases, (g) dehydrogenases, (h) kinases, (i) demethylases, (j) phosphatases, (k) DNA-dependant and (l) RNA-dependant in *L6*-NaCl was compared with *L6*-unntreatd line and represented in the form of heat maps. All these catalases exhibited an expression of >1.3-fold and higher than WT-untreated, WT-NaCl and *L6*-untreated samples.

##### 3.7.1.2. Oxidoreductases

The oxidoreductases constitute about 2% of the total highly expressed catalases. These proteins are reported to be mainly involved in anti-stress processes and ROS homeostasis. NAD(P)H-quinone oxidoreductase subunit M, copper chaperone for superoxide dismutase, superoxide dismutase [Fe] 1, superoxide dismutase [Mn] and peptide methionine sulfoxide reductase A5 were highly expressed (Fig. 4b). NAD(P)H-quinone oxidoreductase (NQOR) is a detoxification enzyme that converts reactive quinones and quinone-imines into less reactive and less toxic hydroquinone forms. This enzyme is also responsible for the oxidation of NADH, a potential source of NAD^+^. The NADH/NAD^+^ ratio is vital in regulating ATP synthesis and other cellular pathways (Melo *et al*., 2004).

##### 3.7.1.3. Lyases

Sphingosine-1-phosphate lyase and two isoforms of phenylalanine ammonia lyases-PAL1 and PAL2 were found to have an expression of 1.78, 1.7 and 1.5 folds, respectively (Fig. 4c). PALs catalyzes the deamination of phenyl alanine from primary metabolism to secondary phenolic metabolism in plants (Hahlbrock & Scheel, 1989).

##### 3.7.1.4. Ligases

About 8% of the total highly expressed catalases were ligases. Four group of ligase proteins such as E3 ubiquitin-protein ligases, coumarate-CoA ligases, cysteine ligase and DNA ligase were expressed in *L6*-NaCl line. The members of E3 ubiquitin-ligases that were expressed included CCNB1IP1 homolog (Cyclin B1 Interacting Protein 1), BRE1-like 1, BRE1-like 2, ATL31, ATL41, ZFP1 (zinc-finger protein 1), XBOs35 and EL5 proteins (Fig. 4d). In addition to these, phospho pantothenate-cysteine ligase 1 and DNA ligase 4 were also expressed at higher levels. BRE1 and BRE2 are yeast homologs of plant histone monoubiquitination 1 (HUB1) and 2 (HUB2) proteins, respectively, which are the cell cycle check point related proteins (Fleury *et al*., 2007). The 4-coumarate-CoA ligase (CCL) proteins exist as multiple isozymes and catalyze the conversion of 4-coumarate to CoA esters in phenylpropanoid metabolism, thereby generating many secondary compounds. Seven isoforms of CCLs *viz*., 4CCL3, 4CCL4, 4CCL5, 4CCL6, 4CCL7, 4CCL8, 4CCL9 were expressed. Each of these isoforms has different substrate affinities indicating their specific roles in plant metabolism (Hamberger & Hahlbrock, 2004).

##### 3.7.1.5. Hydrolases

In plants, the Indole-3-acetic acid (IAA), the abundant form of plant auxins, exists as inactive amide-linked conjugates, which are hydrolyzsed by a family of IAA-amino acid hydrolase 1-like (ILR1-like or ILL) proteins into free and active IAA (Bartel & Fink, 1995). Four ILLs *viz*., ILL2, ILL3, ILL6 and ILL8 were highly expressed in the *L6*-NaCl line (Fig. 4e). Among these, ILL2 was found to have highest catalytic activity with ILR-Ala being its substrate (Carranza *et al*., 2016).

##### 3.7.1.6. Peptidases

Four group of peptidases such as glutamate carboxypeptidase (PLA3), signal peptide peptidase-like 3 (SPPL3), stromal processing peptidase (SPP) and leucine aminopeptidase (LKHA4) were expressed (Fig. 4f). SPP, which showed a 1.3 fold expression is a chloroplast processing endopeptidase (CPE) that removes transit peptides from precursors of chloroplast-targeted proteins involved in photosynthesis (Richter & Lamppa, 1998). SPPL3, which is expressed upto 1.7-fold is a Golgi-localized aspartic proteinase and a member of intramembrane cleaving proteases (I-CLiPs). SPPL3 has been shown to be involved in organogenesis, gametophyte development, pollen maturation (Tamura *et al*., 2008, Han *et al*., 2009, Voss *et al*., 2013). The rice glutamate carboxypeptidase that is expressed upto 1.5-fold is encoded by PLASTOCHRON3 (PLA3) that catalyzes small peptides into signaling molecules regulating multiple physiological functions (Kawakatsu *et al*.,, 2009). Leucine aminopeptidase (LKHA4) and aminopeptidases M1-D were expressed upto 1.4 and 1.7-folds, respectively. These proteins are associated with plant defence responses (Chao *et al*., 1999).

##### 3.7.1.7. Dehydrogenases

Eight different dehydrogenases that are involved in various physiological processes were expressed to higher levels in a slected *L6*-NaCl rice transgenic line. These include 6-phosphogluconate dehydrogenase, decarboxylating 1 (6PGD1), cytokinin dehydrogenase 8 (CKX8), D-2-hydroxyglutarate dehydrogenase (D2HDH), malate dehydrogenase (MDH), betaine aldehyde dehydrogenase 2 (BADH2), cinnamyl alcohol dehydrogenase 8B (CAD8-B), cinnamyl alcohol dehydrogenase 7 (CAD7) and glycerol-3-phosphate dehydrogenase [NAD(+)] 3 (GPDH3) (Fig. 4g).

##### 3.7.1.8. Kinases

Kinases constituted a large percentage (30%) of the total highly expressed catalases. Nearly 63 individual protein kinases were expressed. These include members of ser/thr kinases, calcineurin B-like (CBL)-interacting protein kinases (CIPKs), hexokinases (HXKs), cyclin-dependant kinases (CDKs), histidine kinases (HKs), adenylate kinases (ADKs), calcium-dependant protein kinases (CDPKs), shikimate kinase (SK2), mitogen-activated protein kinases (MAPKs), LRR-receptor kinases (LRR-RK), phytol kinases (PHYK), G-type lectin S-receptor-like serine/threonine-protein kinases, GTP diphosphokinases etc. Eight ser/thr protein kinases were found to be expressed. CBL-CIPKs like CIPK4, CIPK5, CIPK12, CIPK24, CIPK28 and CIPK29 were expressed (Fig. 4h).

The glucose-mediated signaling induces transcriptional activation of thousands of genes involved in a wide range of biological activities. Hexokinases and TOR kinase are two of the three glucose-modulated master regulators in plants with the former acting as a Glc-sensor whereas the presence of Glc activates the latter (Sheen, 2014). Rice HXKs are encoded by a family of ten genes, among which HXK1, HXK6, HXK8 and HXK9 showed high expression in our study. While OsHXK1 is mitochondrial, HXK6 mobilizes between mitochondria and the nucleus and performs dual function like acting as both Glc-sensor and Glc-metabolizing enzyme (Sheen, 2014; Aguilera-Alvarado & Sánchez-Nieto, 2017). NEK3 and NEK6 were expressed to higher levels in *L6*-NaCl line and these are associated with cell cycle regulation, particularly spindle bipolarity during mitosis (Chang *et al*., 2009). ATR kinase, one of the central regulators of DNA damage induced by reactive oxygen intermediates (Maréchal & Zou, 2013) was highly expressed upto 1.6-fold. Four HKs (HK1, HK3, HK4 and HK5) that are cytokinin receptors and regulated by salt stress were also highly expressed. Among the CDKs; CDKE1, CDKG1, CDKC2, CDKD1 were highly expressed.

##### 3.7.1.9. Demethylase

Lysine-specific histone demethylase (JMJ703) was the only demethylase that was highly expressed (Fig. 4i). This lyase either activates or represses transcription, thereby playing an important role in regulation of gene expression in a reversible manner (Anand & Marmorstein, 2007). Jumonji (JMJ) proteins mediate histone demethylation and are found to regulate brassinosteroid signal transduction and floral organ development, and also stress tolerance in plants (Pandey *et al*., 2002; Tsukada *et al*., 2006; Chen *et al*., 2013; Shen *et al*., 2014).

##### 3.7.1.10. Phosphatases

About 14% of the total highly expressed catalases were phosphatases in the present analysis. Protein phosphatases modulate protein phosphorylation by reversing the reactions catalyzed by protein kinases. Rice possesses 78 Mg^2+^-dependant type 2C protein phosphatases (PP2Cs), of which 26 were expressed in this study (Fig. 4j). Protein phosphatase 2A-B (PP2A-B) is a ser/thr phosphatase was found to influence plant development, hormone signaling and salt stress responses (Chen *et al*., 2014). Another highly expressed protein, BSL2 (BRASSINOSTEROID-INSENSITIVE1 SUPPRESSOR 1-like protein 2) is a ser/thr protein phosphatase that is an effector of brassinosteroid signaling pathway playing an important role in determing the grain length (Maselli *et al*., 2014).

##### 3.7.1.11. DNA-mediated catalytic activities

The enzymes involved in DNA replication such as DNA replication licensing factors (MCM2, 6 and 8), replication proteins A-70 (RPA70-A and B), DNA polymerase α-catalytic subunit; gene expression regulation protein-DNA (cytosine-5)-methyltransferase 1A, DNA repair like ATP-dependent DNA helicase 2 (KU80) and RAD54 (DNA repair and recombination protein) were also expressed in the slected rice *L6*-NaCl line (Fig. 4k).

##### 3.7.1.12. RNA-mediated catalytic activities

DEAD-box ATP-dependant RNA helicases (DEAD-RHs) have roles in many cellular activities including RNA processing, nuclear export of RNA, ribosome assembly and translation, and are also involved in transcription regulation by acting as coactivators or corepressors of transcription factors (Liu & Imai, 2018). Given their importance in ribosome biogenesis, the high expression of as many as 19 DEAD-RHs in the present study can be correlated. The S-adenosyl-L-methionine-dependent tRNA 4-demethylwyosine synthase and wybutosine-synthesizing protein 2/3/4 were also found to be highly expressed (Fig. 4l). These proteins modify the guanosine residues present adjacent to the anticodon of phenylalanine tRNA (Noma & Suzuki, 2006). This modification stabilizes the codon-anticodon interactions during decoding on the ribosome, thereby ensuring accurate protein translation.

#### 3.7.2. Transporters

After catalases, transporters occupy a major portion of the HEPs (20%). About 66 different transporters were expressed which included 13 members of ATP-binding cassette G-family (ABC-G) proteins and 30 ion transporters. Of the ion transporters, nine of them were different homologues of potassium transporters (HAKs) (Fig. 5a). The ABC-G transporters drive the intercellular exchange of phytohormones and secondary metabolites, thus playing an important role in various physiological functions (Hwang *et al*., 2016). ABC-G13, 35, 37, 38, 40, 42, 44, 45, 46, 48, 49, 52 and 53 were expressed in the present analysis. Because plants synthesize different metabolites, they contain a large number of these transporters whose expression would be tissue/substrate/condition specific.

**Figure 5.**
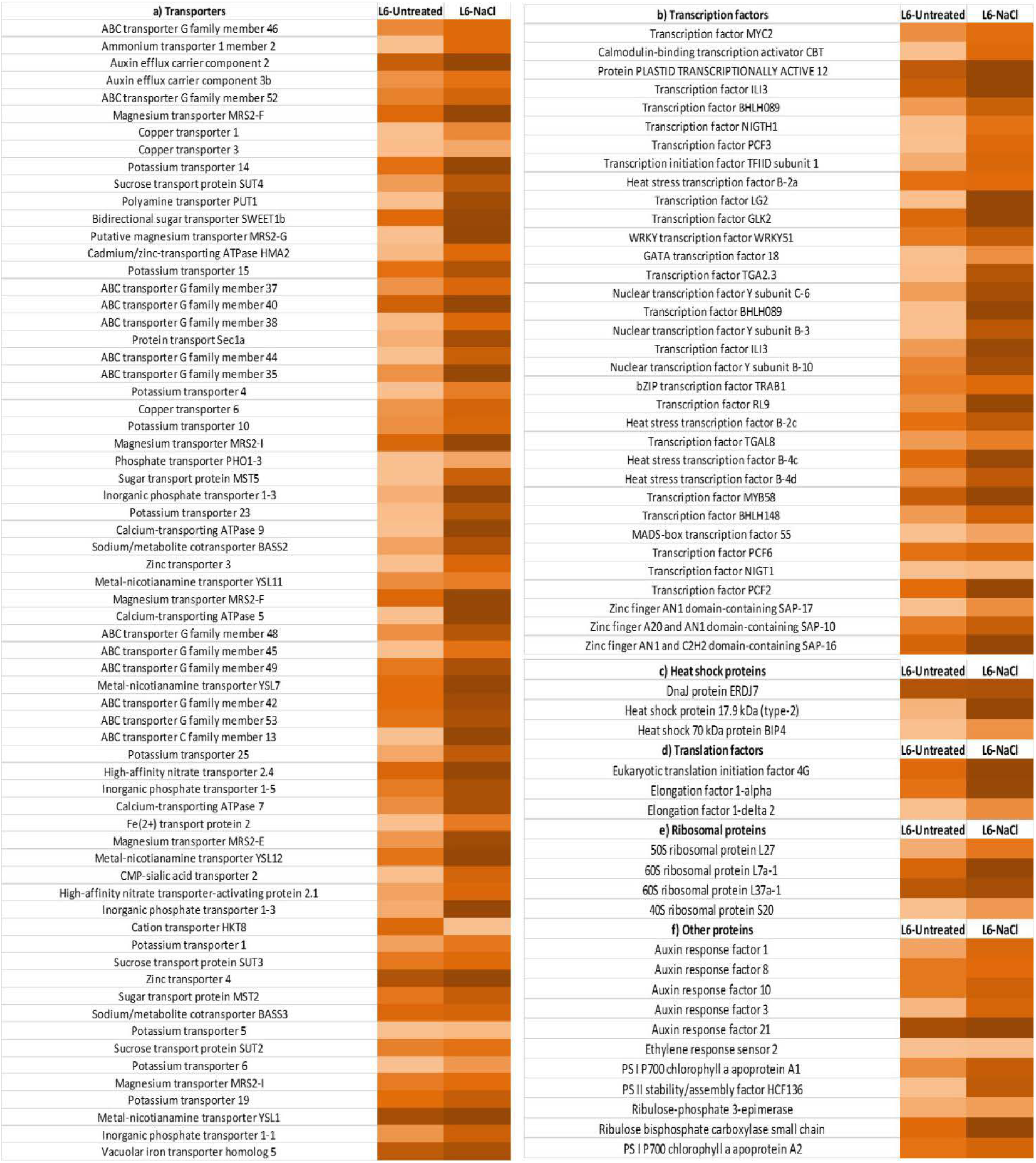
Expression pattern of transporters, transcription factors and translation-related proteins. The level of expression of (a) transporter, (b) transcription factors, (c) heat shock proteins, (d) translation factors, (e) ribosomal and (f) other highly expressed proteins were depicted in the form of heat maps between *L6*-NaCl and *L6*-unntreatd lines. All these proteins exhibited an expression of >1.3-fold and higher than WT-untreated, WT-NaCl and *L6*-untreated samples.

#### 3.7.3. Transcription factors

About 34 transcription factors (TFs) occupying 10% of the total HEPs in *L6*-NaCl line are involved in developmental, hormone and stress signalling pathways (Fig. 5b). Among these, transcription factor ILI3 (TFILI3), Heat stress transcription factor B-4c (HSFB4C), TF-MYB58, TF-PCF2, stress-associated protein 10 (SAP10) and 16 (SAP16) were highly expressed. HSFs are activated by phosphorylation mediated by MAPKs (MAPK6), which in turn bind to heat stress elements (HSEs) and elicit the expression of stress responsive genes (Pérez-Salamó *et al*., 2014; Guo et al, 2016). The other TFs that are expressed such as WRKY and MYB are also activated by MAPK6 (Li *et al*., 2012). Nuclear Factor Y subfamily-C (NFY-C6) is a novel histone-like TF that was also highly expressed (1.93-fold).

#### 3.7.4. Heat shock proteins

The heat shock proteins that were expressed include HSP2, HSP70-BIP4 and HSP40-DNAJ7 (ERDJ7) (Fig. 5c). HSPs are molecular chaperones that trigger the defence-related unfolded response pathway (UPR) against adverse environmental conditions. BIP4 and ERDJs are components of UPR that serves to inhibit improper protein translation of nascent polypeptides or degradation of misfolded proteins, thereby functionig as a protein checking machinery (Ohta *et al*., 2013; Ohta & Takaiwa, 2014). Although the role of OsERDJ7 is not characterized, its expression in salt tolerant line might provide a clue that it is also associated in stress responses as other J proteins.

#### 3.7.5. Translation-related proteins

The translation factors, translation initiation factor 4G, elongation factor 1-alpha and 1-delta and, ribosomal proteins like RPL7, RPL37a-1 and RPS20 were also expressed to higher leves (Fig. 5d).

#### 3.7.6. Expression of other proteins

The other group of proteins that were highly expressed include Auxin response factors (ARF1, 3, 8, 10 and 21), ethylene response sensor 2 (ERS2) and photosystem stabilizing proteins (PS-I P700 chlorophyll a apoprotein A1, A2 and PS-II stability/assembly factor HCF136) (Fig. 5e & f).

### 3.8. Validation of protein expression by qRT-PCR

We validated the transcript levels of some of the randomly selected genes whose proteins were highly expressed in iTRAQ technique. These include TOR, WRKY51, RPL7, RPL37 and RPS20, which were expressed by 1.9, 1.7, 2, 1.95 and 1.42 protein folds, respectively. After 200 mM NaCl treatment, the transcripts of all the selected genes were significantly up-regulated with *OsTOR, OsRPL37* and *OsRPL7* showing >15-fold up-regulation compared with their respective controls (Supplementary Fig. 6). The transcript levels of *RPL7* was found to be highest (32-fold). These results supports the expression pattern of proteins obtained by iTRAQ.

### 3.9. Network analysis of HEPs

The association network of HEPs and RPL6 was constructed by submitting the candidate proteins in the STRING, which builds networks based on functional association (indirect interactions) and also includes direct physical interactions. Out of 333 submitted proteins, the database identified 300 proteins and the interaction analyses were shown for these 300 protein nodes having 472 edges with 3.15 average node degree, 0.403 average local clustering coefficient and a PPI enrichment p-value of 2.39e-11. The higher number of edges (472) compared to the expected number (343) indicates significantly higher interactions or connections (Fig. 6). Such an enrichment also suggests that the proteins are at least partially biologically connected, as a group. Based on these interactions, we tried to link the pathways that these proteins are a part of and are responsible for growth and tolerance under salt stress (Fig. 7). The RPL6 appears to interact with five nodes; elongation factor 1-alpha, RPS20, RPL37, RPL7 and DEAD-box ATP-dependent RNA helicase 47B. However, the reaction (transcription/translation/catalysis) that results from these bindings needs to be investigated further. The functional networking of each of these nodes triggers the activities of downstream targets, which together promote growth and tolerance in response to salt stress.

**Figure 6.**
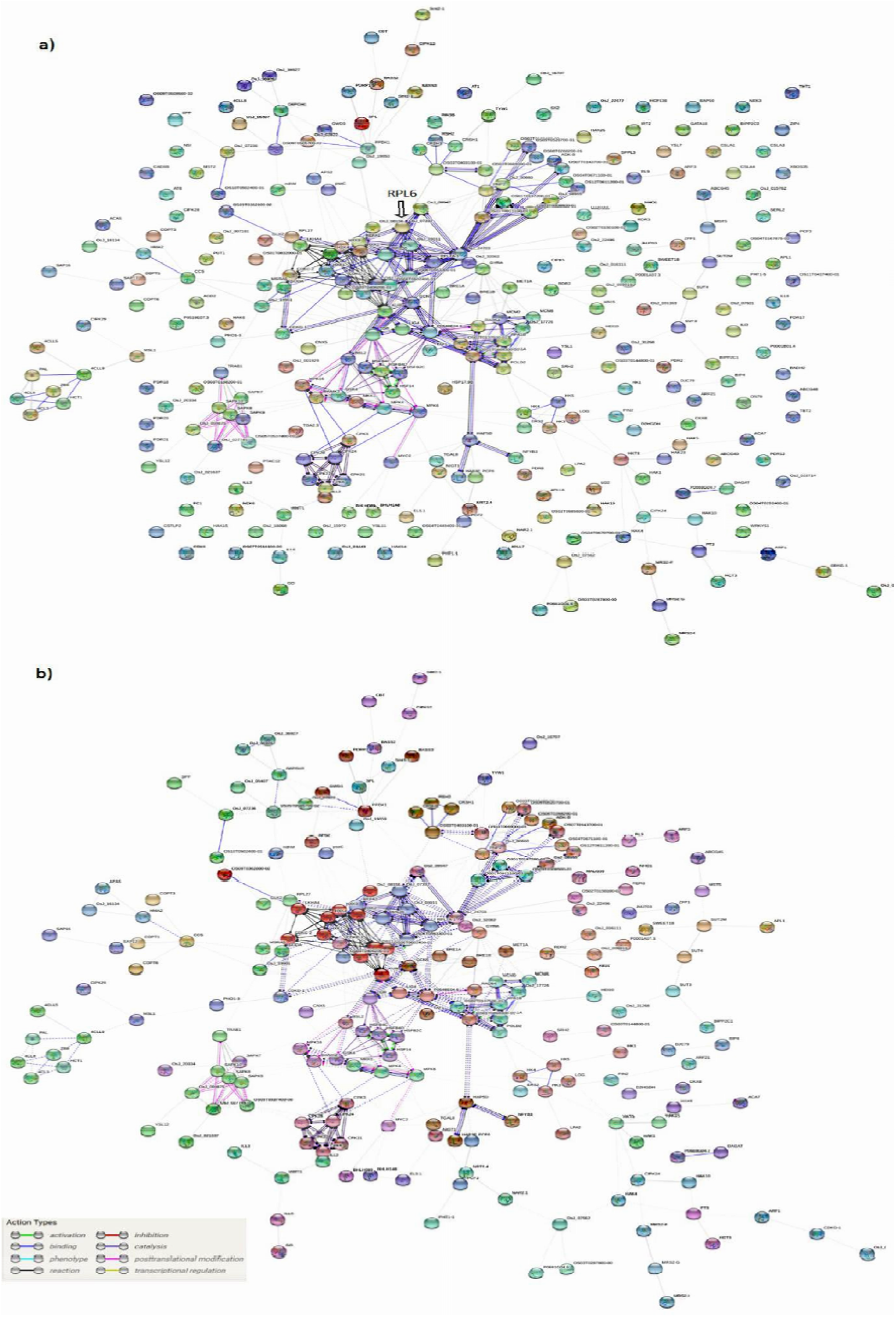
*In silico* association networks of highly expressed proteins. The network analysis of (a) 333 highly expressed proteins were performed in STRING database, of which approximately 50 nodes were not part of this interaction.. The protein nodes that were part of the network were (b) MCL-clustered (Markov Cluster algorithm) with an inflation parameter of three. The types of node interaction was depicted with different colors which is provided at the bottom left corner.

**Figure 7.**
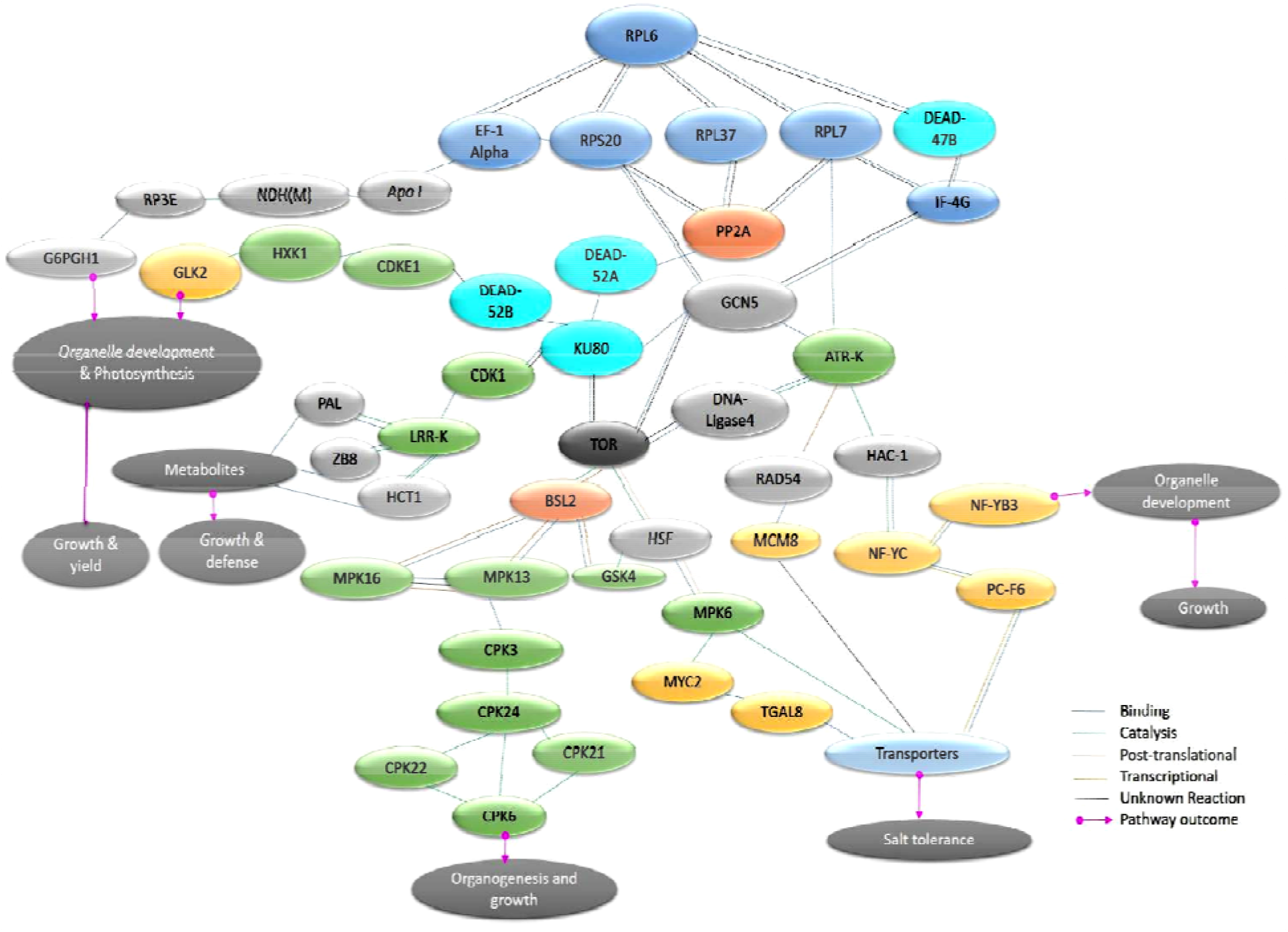
Overview of networking of highly expressed proteins in growth and salt tolerance. Based on the protein interaction network obtained from STRING, a pathway is predicted involving the highly expressed proteins that resulted in growth and tolerance under salt stress conditons.

## 4. DISCUSSION

Rice being a glycophyte is susceptible to high sodium levels, which adversely affect seedling, vegetative and reproductive stages, wrecking its growth and productivity. Therefore, it is important to comprehend the physiological processes occurring during salt stress so as to develop salt-tolerant rice by transgenic and genome editing strategies or suitable agronomic practices. When plants perceive stress signals, a cascade of signaling events is initiated leading to physiological and metabolic changes ensuring the survival of plant under adverse conditions. Identification of novel candidate genes and alleles for enhanced tolerance to salinity and also for maintaining stable yield under stress is of paramount importance in this context (Koyama *et al*., 2001; Hu *et al*., 2008; Ye *et al*., 2009). Ribosomal proteins are known to play an integral role in generation of rRNA structure and forming protein synthesizing machinery in cells (Rodnina & Wintermeyer, 2009). They are also crucial for growth and development of all organisms (Ishii *et al*., 2006). Since salt stress can result in modification of protein synthesis, it has been observed in many instances that upregulation of genes encoding ribosomal proteins in plants under stressed condition led to efficient reconstruction of protein synthesizing machinery in cells.

### 4.1. Salt tolerance phenomenon in *L6* expressing rice transgenic plants

We have reported the involvement of *RPL6* in enhancing WUE in rice earlier in our gain-of-function mutagenesis studies in rice cultivar BPT5204 (Moin *et al*., 2016a). In the present study, we show that the transgenic rice plants overexpressing *RPL6* were also found to be tolerant to moderate (150 mM) to high (200 mM) salt concentrations at the seedling stage. When these were shifted to salt-free medium, some of the seedlings from high expression line not only revived but also exhibited growth and yield parameters nearly equivalent to the WT grown without NaCl stress. The salt tolerant high expression lines also showed high chlorophyll contents, quantum efficiency and accumulated higher quantities of osmolyte, proline. Our overall findings showed that high expression lines of *L6* conferred NaCl tolerance in transgenic rice without much compromise on growth and productivity. These results prompted us to investigate the complete protein profile of a selected high expression *L6*-NaCl line. In view of this, an integrated proteomic approach, iTRAQ (Isobaric tags for relative and absolute quantitation) combined with high-throughput mass spectrometry (LC-MS/MS) was employed to identify the key proteins that were particularly highly expressed in the *L6-5* transgenic line after 200 mM NaCl stress treatment. The identification of these proteins, their network and the pathways they mediate would help us understand the mechanism of *L6*-mediated NaCl tolerance. A wide variety of proteins such as those involved in growth and developmental processes, immune responses, cellular homeostasis, signal transduction (transcription and translation), membrane and organelle transport, binding and catalytic activities were identified.

### 4.2. Involvement of various highly expressed proteins in plant growth, and salt stress tolerance

All the oxidoreductases that were highly expressed were found to be important members of photosynthetic and ROS scavenging system, which participate in many reactive oxygen metabolism related processes in cells and are responsible for maintaining normal cell metabolism. The ROS-scavenging proteins were also shown to be involved in improving growth and development of rice under abiotic conditions including salt stress (Zhang *et al*., 2013). The PAL1 and PAL2 proteins that were highly expressed in the present study are involved in the biosynthesis of phytohormones, phenylpropanoids, cuticular wax and play important functions in photosystem stability and defense in response to abiotic and biotic challenges (Kumar & Ellis, 2001). The reduced function of PAL proteins was found to have a negative impact on disease resistance and tolerance to abiotic stresses (Cass *et al*., 2015). The enzymatic activity of PAL was found to be increased after salt treatment along with other redox enzymes that is indicative of its role in salt-induced antioxidative mechanism (Gholizadeh & Kohnehrouz, 2010). A large set of E3 ubiquitin and coumarate-Co A ligases (CCLs) were also highly expressed in our study. The ligase proteins such as HUB, CCNB1IP, ZFP and EL RING-type ligases that were highly expressed in the present analysis were reported to be involved in cell cycle regulation and meotic crossover, which play important roles in organogenesis and gametogenesis and hence in leaf, shoot, root and floral morphogenesis (Chrispeels *et al*., 2001; Koiwai *et al*., 2007; Nishizawa *et al*., 2008; Maekawa *et al*., 2012; Wang *et al*., 2012). The high expression of these cell cycle regulators and check point proteins ensures that the process of cytokinesis takes place precisely in an orderly manner under the conditions of stress also. The CCLs synthesize metabolites such as lignin, anthocyanins, chalcones, aurones, isoflavonoids, furanocoumarins, flavones and flavonols, which have diverse functions in providing rigidity to the plant, pigmentation and protection against environmental cues (Hamberger & Hahlbrock, 2004). Overexpression of some of these CCLs has also been shown to result in transgenic plants with reduced accumulation of ROS and increased tolerance to osmotic stresses (Chen *et al*., 2019).

The *L6*-NaCl line also showed the high expression of cytokinin riboside 5’□ monophosphate phosphoribohydrolase (CR5MPRH), an important enzyme of the single□step cytokinin activation pathway that is encoded by one of the seven LONELY GUY (LOG) homologues in rice (Kurakawa *et al*., 2007). Cytokinins, whose activation and spatiotemporal distribution are finely regulated by enzymes, play vital roles in processes such as root proliferation, apical dominance and phyllotaxis (Shimizu-Sato *et al*., 2009). Cytokinins are also the inducers of cytokinesis in the presence of auxins (Mok & Mok, 2001). Four ILL isoforms that are involved in generating active form of IAA were highly expressed in our study. These have been directly linked with growth, development and abiotic stress tolerance with overexpression of ILL3 rendering transgenic plants higher salt tolerance (Junghans *et al*., 2006). The coordinated high expressin of auxin and cytokinin activating enzymes ensures that the cross-linked biological processes mediated by these phytohormones are not hampered under salt stress conditions in *L6* transgenic plants. The 6PGD1 of oxidative pentose phosphate pathway, MDH, GPDH3 and CAD are important regulatory enzymes that were also highly expressed in the present analysis. These enzymes functions in maintaining redox homeostasis and are mainly evolved to provide tolerance against oxidative stress damage (Shen *et al*., 2006; Hebbelmann *et al*., 2012; Esposito, 2016; Kim *et al*., 2019).

The *L6*-NaCl line also exhibited high expression of histidine kinases and HSPs. Arabidopsis *ahk* mutants showed low proline accumulation along with high salt sensitivity (Kumar & Verslues, 2015). Plants overexpressing members of UPR system were found to have stable photosystems, absorb less Na^+^ and accumulate more osmolytes under salt stress resulting in salt tolerance (Fu *et al*., 2016). These studies corroborated our current findings with high expression of HKs and HSPs accompanied with more accumulation of proline, high quantum efficiency and tolerance to salt stress. Simultaneous expression of kinases and type 2C-phosphatases, which are positive and negative regulators of stress signaling pathways, respectively (Schweighofer *et al*., 2004; Xue *et al*., 2008) indicates that RPL6 is involved in balancing of the expression of these multifunctional proteins under stress without compromising the growth and development of *L6* transgenic rice plants. A large group of potassium (HAK) and ABC-G transporters that are involved in the enhancement of crop yield and stress tolerance (Moon & Jung, 2014) were also highly expressed in our study.

A large set of DEAD box RNA helicases involved in ribosome biogenesis through rRNA processing were also highly expressed in *L6*-NaCl line. This process was shown to be controlled by TOR kinase (Xing *et al*., 2019). The TOR is a multifaceted protein kinase involved in modulation of translation, ribosome biogenesis, nutrient and energy signaling, thereby tightly regulating plant growth and development. In addition, TOR has also been shown to play a pivotal role in abiotic stress tolerance, enhancement of water-use efficiency and yield-related traits possibly by up-regulating TOR-complex components (Raptor and LST8) and other stress-tolerant genes in rice (Bakshi *et al*., 2017; Bakshi *et al*., 2018). Some of the CDKs that were highly expressed in this study were found to regulate transcription of downstream genes that encode proteins involved in cell cycle and stress responses (Xiang *et al*., 2007; Ng *et al*., 2013; Zheng *et al*., 2014; Takatsuka *et al*., 2015; Zhao *et al*., 2017). Also, the high expression of ATRK as seen in *L6*-NaCl line might be to ensure the activation of DNA repair pathway against the damage caused by Na^+^-induced ROS during salt stress. The high expression of other kinases like SK2, LRR-RK, CBL-CIPKs, CDPKs (CDPK3, 6, 22, 24, 26 and 28), MAPKs (MAPK1, 4, 6, 13 and 16), PHYKs (PHYK1 and 2) and SAPKs (7, 8, 9 and 10) as observed in the current study were found to be common defence responses of plants to abiotic stresses (De Lorenzo *et al*., 2009; Cristina *et al*., 2010; Asano *et al*., 2012; Nongpiur *et al*., 2012; Tohge *et al*., 2013; Basu & Roychoudhury, 2014; Mohanta & Sinha, 2016; Spicher *et al*., 2017; Shi *et al*., 2018). Some of the highly expressed transcription factors in *L6*-NaCl line like TFILI3, HSFB4C, MYB58, TF-PCF2, SAP10 and SAP16 are also involved in the regulation of environmental stress responses with the overexpression of several of them was found to have conferred tolerance to salt and other aiotic stresses (Mukhopadhyay *et al*., 2004; Hozain *et al*., 2012; Dixit *et al*., 2018).

### 4.3. *In silico* signaling network analysis and possible mechanism of salt tolerance in *L6* expressing rice transgenic plants

Our protein-protein network analysis showed that TOR might activate the HSF possibly by its phosphorylation. This HSF binds with MAPK6 and acts in a feedback regulatory loop (Pérez-Salamó *et al*., 2014) activaing other transcription factors like MYC2 and TGAL8, which further affects the activities of many transporter proteins. The MAPK6, which was highly expressed in *L6*-NaCl line mediates the regulation of various biotic and abiotic defense responses by coordinating the activity of transcription factors which, in turn, control the targeted expression of a large group of genes (Pérez-Salamó *et al*., 2014). The protein network analysis also predicted the TOR-mediated regulation of calcium-dependant protein kinases (CPK3, 24, 22, 21 and 6) via MAPK16 and MAPK13. This circuit functions in signal transduction pathways, positively regulating responses to ABA, which in turn is involved in regulating growth and stress responses. This coordinated expression of TOR, HSF and MAPKs and their downstream transcription factors in our study might have been one of the important reasons for salt tolerance. NFY-C, which was also activated in our activation tagged mutants screened for salt tolerance (Manimaran *et al*., 2017), appears to interact with transcription factor, PCF6 that further regulates the activities of different ion transporters. The hexokinases, DEAD-box helicases and GLK2 transcription factor were other major group of networking proteins playing pivotal roles in organogenesis, chlorophyll biosynthesis, photosynthesis and plant defense.

A general phenomenon involving salt tolerance occurs either by regulating Na^+^ influx into cells or intracellular sequestration. While sodium compartmentation into vacuoles is mediated by Na^+^/H^+^ antiporter encoded by SOS1, its influx is regulated by importantly by potassium trasnporters (HAKs, Wu, 2018). Some of these transporters including those that expressed in the current study like OsHAK1, OsHAK5 and OsHAK6 compete with Na^+^ for K^+^ under high salt conditions or potassium starvation to maintain cellular ionic homeostasis (Horie *et al*., 2011; Chen *et al*., 2015). Expression of specific members of HAK transporters might be an important reason for tolerance against salt stress in certain wild type rice genotypes that are being used as potential donors for improving salt tolerance in other rice cultivars (Quan *et al*., 2018). Expression of these transporters in *L6*-NaCl indicates that RPL6 induces salt tolerance possibly by restricting sodium entry by expressing Na^+^ insensitive K^+^ transporters instead of its sequestration as there was no expression of Na^+^/H^+^ antiporter, which has been reported to be generally associated with salt tolerance in other studies (El Mahi *et al*., 2019). The networking of each one of the highly expressed proteins in a functional circuit might be responsible for promoting the growth and yield under salt stress conditions in *RPL6* transgenic rice.

## Author contribution statement

MM, PBK and MSM designed the experiments and MM performed all the experiments. AS and AB helped in salt stress screening and qRT-PCR experiments. MM and PBK prepared the manuscript. All the authors read and approved the manuscript.

## Acknowledgements

This investigation forms a part of the INSPIRE-faculty project on abiotic stress responsiveness of ribosomal large subunit proteins funded by the Department of Science and Technology (DST), Government of India through grant number IFA17-LSPA67 to MM. MM acknowledges the Fellowship and Research grants received through this DST-INSPIRE faculty program. Authors also acknowledge the facilities obtained from the Department of Biotechnology, ICAR-Indian Institute of Rice Research (IIRR), Hyderabad.

## Conflict of interest

The authors declare the absence of any commercial or financial interests that could be constructed as potential conflicts of interest.

## Abbreviations

RP: Ribosomal Protein
RPL: Ribosomal Protein Large subunit
RPS: Ribosomal Protein Small subunit
DEP: differentially expressed proteins

## Notes

### Competing Interest Statement

The authors have declared no competing interest.

## References

Aguilera-Alvarado, G. P., & Sánchez-Nieto, S. (2017) Plant hexokinases are multifaceted proteins. Plant and Cell Physiology, 58, 1151–1160.

Anand, R., & Marmorstein, R. (2007) Structure and mechanism of lysine-specific demethylase enzymes. Journal of Biological Chemistry, 282, 35425–35429.

Asano, T., Hayashi, N., Kikuchi, S., & Ohsugi, R. (2012) CDPK-mediated abiotic stress signaling. Plant Signaling & Behavior, 7, 817–821.

Ashraf, M. F. M. R., & Foolad, M. (2007) Roles of glycine betaine and proline in improving plant abiotic stress resistance. Environmental and experimental bot. 59, 206–216.

Bai, D., Zhang, J., Xiao, W., & Zheng, X. (2014) Regulation of the HDM2-p53 pathway by ribosomal protein L6 in response to ribosomal stress. Nucleic Acids Research, 42, 1799–1811.

Bakshi, A., Moin, M., Datla, R., & Kirti, P. B. (2017) Expression profiling of development related genes in rice plants ectopically expressing AtTOR. Plant Signaling & Behavior, 12, e1362519.

Bakshi, A., Moin, M., Kumar, M. U., Reddy, A. B. M., Ren, M., Datla, R., Siddiq, E.A., & Kirti, P.B. (2017) Ectopic expression of Arabidopsis Target of Rapamycin (AtTOR) improves water-use efficiency and yield potential in rice. Scientific Reports, 7, 42835.

Bartel, B., & Fink, G. R. (1995) ILR1, an amidohydrolase that releases active indole-3-acetic acid from conjugates. Science, 268, 1745–1748.

Basu, S., & Roychoudhury, A. (2014) Expression profiling of abiotic stress-inducible genes in response to multiple stresses in rice (Oryza sativa L.) varieties with contrasting level of stress tolerance. BioMed Research International, 2014.

Bohnert, H. J., Ayoubi, P., Borchert, C., Bressan, R. A., Burnap, R. L., Cushman, J. C., … & Hasegawa, P. M. (2001) A genomics approach towards salt stress tolerance. Plant Physiology and Biochemistry, 39, 295–311.

Carranza, A. P. S., Singh, A., Steinberger, K., Panigrahi, K., Palme, K., Dovzhenko, A., & Dal Bosco, C. (2016) Hydrolases of the ILR1-like family of Arabidopsis thaliana modulate auxin response by regulating auxin homeostasis in the endoplasmic reticulum. Scientific Reports, 6, 24212.

Cass, C. L., Peraldi, A., Dowd, P. F., Mottiar, Y., Santoro, N., Karlen, S. D., & Moskvin, O. V. (2015) Effects of PHENYLALANINE AMMONIA LYASE (PAL) knockdown on cell wall composition, biomass digestibility, and biotic and abiotic stress responses in Brachypodium. Journal of Experimental Bot. 66, 4317–4335.

Chang, J., Baloh, R. H., & Milbrandt, J. (2009) The NIMA-family kinase Nek3 regulates microtubule acetylation in neurons. Journal of Cell Science, 122, 2274–2282.

Chao, W. S., Gu, Y. Q., Pautot, V., Bray, E. A., & Walling, L. L. (1999) Leucine aminopeptidase RNAs, proteins, and activities increase in response to water deficit, salinity, and the wound signals systemin, methyl jasmonate, and abscisic acid. Plant Physiology, 120, 979–992.

Chapman, J. R., Barral, P., Vannier, J. B., Borel, V., Steger, M., Tomas-Loba, A., et al. (2013) RIF1 is essential for 53BP1-dependent nonhomologous end joining and suppression of DNA double-strand break resection. Molecular Cell, 49, 858–871.

Chen, G., Hu, Q., Luo, L. E., Yang, T., Zhang, S., Hu, Y., et al. (2015) Rice potassium transporter O s HAK 1 is essential for maintaining potassium mediated growth and functions in salt tolerance over low and high potassium concentration ranges. Plant, Cell & Environment, 38, 2747–2765.

Chen, J., Hu, R., Zhu, Y., Shen, G., & Zhang, H. (2014) Arabidopsis PHOSPHOTYROSYL PHOSPHATASE ACTIVATOR is essential for PROTEIN PHOSPHATASE 2A holoenzyme assembly and plays important roles in hormone signaling, salt stress response, and plant development. Plant Physiol. 166, 1519–1534.

Chen, L., Zhao, J., Song, J., & Jameson, P. E. (2020) Cytokinin dehydrogenase: a genetic target for yield improvement in wheat. Plant Biotechnology Journal, 18, 614–630.

Chen, Q., Chen, X., Wang, Q., Zhang, F., Lou, Z., Zhang, Q., & Zhou, D. X. (2013) Structural basis of a histone H3 lysine 4 demethylase required for stem elongation in rice. PLoS Genetics, 9(1).

Chen, X., Wang, H., Li, X., Ma, K., Zhan, Y., & Zeng, F. (2019) Molecular cloning and functional analysis of 4-Coumarate: CoA ligase 4 (4CL-like 1) from Fraxinus mandshurica and its role in abiotic stress tolerance and cell wall synthesis. BMC Plant Biology, 19, 231.

Chen, X., Wang, Y., Lv, B., Li, J., Luo, L., Lu, S., et al. (2014) The NAC family transcription factor OsNAP confers abiotic stress response through the ABA pathway. Plant and Cell Physiology, 55, 604–619.

Chrispeels, H. E., Oettinger, H., Janvier, N., & Tague, B. W. (2000) AtZFP1, encoding Arabidopsis thaliana C2H2 zinc-finger protein 1, is expressed downstream of photomorphogenic activation. Plant Mol. Biol. 42, 279–290.

Cristina, M. S., Petersen, M., & Mundy, J. (2010) Mitogen-activated protein kinase signaling in plants. Annual Review of Plant Biology, 61, 621–649.

Dai Yin, C. H. A. O., Luo, Y. H., Min, S. H. I., Da, L. U. O., & Lin, H. X. (2005) Salt-responsive genes in rice revealed by cDNA microarray analysis. Cell Research, 15, 796–810.

De Lorenzo, L., Merchan, F., Laporte, P., Thompson, R., Clarke, J., Sousa, C., & Crespi, M. (2009) A novel plant leucine-rich repeat receptor kinase regulates the response of Medicago truncatula roots to salt stress. The Plant Cell, 21, 668–680.

Degenhardt, R. F., & Bonham-Smith, P. C. (2008) Arabidopsis ribosomal proteins RPL23aA and RPL23aB are differentially targeted to the nucleolus and are disparately required for normal development. Plant Physiology 147, 128–142.

Del Rio, D., Stewart, A. J., & Pellegrini, N. (2005) A review of recent studies on malondialdehyde as toxic molecule and biological marker of oxidative stress. Nutrition, metabolism and cardiovascular diseases, 15, 316–328.

Dixit, A., Tomar, P., Vaine, E., Abdullah, H., Hazen, S., & Dhankher, O. P. (2018) A stress associated protein, AtSAP13, from Arabidopsis thaliana provides tolerance to multiple abiotic stresses. Plant, Cell & Environment, 41, 1171–1185.

El Mahi, H., Pérez-Hormaeche, J., De Luca, A., Villalta, I., Espartero, J., Gámez-Arjona, F., et al. (2019). A critical role of sodium flux via the plasma membrane Na+/H+ exchanger SOS1 in the salt tolerance of rice. Plant Physiology, 180, 1046–1065.

Esposito, S. (2016) Nitrogen assimilation, abiotic stress and glucose 6-phosphate dehydrogenase: The full circle of reductants. Plants, 5, 24.

Fatehi, F., Hosseinzadeh, A., Alizadeh, H., Brimavandi, T., & Struik, P. C. (2012) The proteome response of salt-resistant and salt-sensitive barley genotypes to long-term salinity stress. Molecular Biology Reports, 39, 6387–6397.

Fleury, D., Himanen, K., Cnops, G., Nelissen, H., Boccardi, T. M., Maere, S., … & Micol, J. L. (2007) The Arabidopsis thaliana homolog of yeast BRE1 has a function in cell cycle regulation during early leaf and root growth. The Plant Cell, 19, 417–432.

Fradet-Turcotte, A., Canny, M. D., Escribano-Díaz, C., Orthwein, A., Leung, C. C., Huang, H., et al. (2013) 53BP1 is a reader of the DNA-damage-induced H2A Lys 15 ubiquitin mark. Nature, 499, 50–54.

Fu, C., Liu, X. X., Yang, W. W., Zhao, C. M., & Liu, J. (2016) Enhanced salt tolerance in tomato plants constitutively expressing heat-shock protein in the endoplasmic reticulum. Genet. Mol. Res, 15(2).

Gholizadeh, A., & Kohnehrouz, B. B. (2010) Activation of phenylalanine ammonia lyase as a key component of the antioxidative system of salt-challenged maize leaves. Brazilian Journal of Plant Physiol. 22, 217–223.

Giri, J., Dansana, P. K., Kothari, K. S., Sharma, G., Vij, S., & Tyagi, A. K. (2013) SAPs as novel regulators of abiotic stress response in plants. Bioessays, 35, 639–648.

Guo, M., Liu, J. H., Ma, X., Luo, D. X., Gong, Z. H., & Lu, M. H. (2016) The plant heat stress transcription factors (HSFs): structure, regulation, and function in response to abiotic stresses. Frontiers in Plant Science, 7, 114.

Hahlbrock, K., & Scheel, D. (1989) Physiology and molecular biology of phenylpropanoid metabolism. Annual Review of Plant Biol. 40, 347–369.

Hamberger, B., & Hahlbrock, K. (2004) The 4-coumarate: CoA ligase gene family in Arabidopsis thaliana comprises one rare, sinapate-activating and three commonly occurring isoenzymes. Proceedings of the National Academy of Sciences, 101, 2209–2214.

Han, S., Green, L., & Schnell, D. J. (2009) The signal peptide peptidase is required for pollen function in Arabidopsis. Plant Physiol. 149, 1289–1301.

Hare, P. D., & Cress, W. A. (1997) Metabolic implications of stress-induced proline accumulation in plants. Plant Growth Regulation, 21, 79–102.

Hebbelmann, I., Selinski, J., Wehmeyer, C., Goss, T., Voss, I., Mulo, P., et al. (2012) Multiple strategies to prevent oxidative stress in Arabidopsis plants lacking the malate valve enzyme NADP-malate dehydrogenase. Journal of Experimental Botany, 63, 1445–1459.

Horie, T., Sugawara, M., Okada, T., Taira, K., Kaothien-Nakayama, P., Katsuhara, M., et al. (2011) Rice sodium-insensitive potassium transporter, OsHAK5, confers increased salt tolerance in tobacco BY2 cells. Journal of Bioscience and Bioengineering, 111, 346–356.

Hu, H., Dai, M., Yao, J., Xiao, B., Li, X., Zhang, Q., & Xiong, L. (2006) Overexpressing a NAM, ATAF, and CUC (NAC) transcription factor enhances drought resistance and salt tolerance in rice. Proceedings of the National Academy of Sciences, 103, 12987–12992.

Hwang, J. U., Song, W. Y., Hong, D., Ko, D., Yamaoka, Y., Jang, S., et al. (2016) Plant ABC transporters enable many unique aspects of a terrestrial plant’s lifestyle. Molecular Plant, 9, 338–355.

Ishii, K., Washio, T., Uechi, T., Yoshihama, M., Kenmochi, N., & Tomita, M. (2006) Characteristics and clustering of human ribosomal protein genes. BMC Genomics, 7, 37.

James, R. A., Blake, C., Byrt, C. S., & Munns, R. (2011). Major genes for Na+ exclusion, Nax1 and Nax2 (wheat HKT1; 4 and HKT1; 5), decrease Na+ accumulation in bread wheat leaves under saline and waterlogged conditions. Journal of Exp. Bot. 62, 2939–2947.

Jiang, D., Zhou, L., Chen, W., Ye, N., Xia, J., & Zhuang, C. (2019) Overexpression of a microRNA-targeted NAC transcription factor improves drought and salt tolerance in Rice via ABA-mediated pathways. Rice, 12, 76.

Junghans, U., Polle, A., Düchting, P., Weiler, E., Kuhlman, B., Gruber, F., & Teichmann, T. (2006) Adaptation to high salinity in poplar involves changes in xylem anatomy and auxin physiology. Plant, Cell & Environ. 29, 1519–1531.

Kawakatsu, T., Taramino, G., Itoh, J. I., Allen, J., Sato, Y., Hong, S. K., et al. (2009) PLASTOCHRON3/GOLIATH encodes a glutamate carboxypeptidase required for proper development in rice. The Plant Journal, 58, 1028–1040.

Kim, Y. H., & Huh, G. H. (2019) Overexpression of cinnamyl alcohol dehydrogenase gene from sweetpotato enhances oxidative stress tolerance in transgenic Arabidopsis. In Vitro Cellular & Developmental Biology-Plant, 55, 172–179.

Koiwai, H., Tagiri, A., Katoh, S., Katoh, E., Ichikawa, H., Minami, E., & Nishizawa, Y. (2007) RING H2 type ubiquitin ligase EL5 is involved in root development through the maintenance of cell viability in rice. The Plant Journal, 51, 92–104.

Kumar, A., & Ellis, B. E. (2001) The phenylalanine ammonia-lyase gene family in raspberry. Structure, expression, and evolution. Plant Physiol. 127, 230–239.

Kumar, M. N., & Verslues, P. E. (2015) Stress physiology functions of the Arabidopsis histidine kinase cytokinin receptors. Physiologia Plantarum, 154, 369–380.

Kurakawa, T., Ueda, N., Maekawa, M., Kobayashi, K., Kojima, M., Nagato, Y., et al. (2007) Direct control of shoot meristem activity by a cytokinin-activating enzyme. Nature, 445, 652–655.

Lee, D. K., Chung, P. J., Jeong, J. S., Jang, G., Bang, S. W., Jung, H., et al. (2017) The rice Os NAC 6 transcription factor orchestrates multiple molecular mechanisms involving root structural adaptions and nicotianamine biosynthesis for drought tolerance. Plant Biotechnology Journal, 15, 754–764.

Li, G., Meng, X., Wang, R., Mao, G., Han, L., Liu, Y., & Zhang, S. (2012) Dual-level regulation of ACC synthase activity by MPK3/MPK6 cascade and its downstream WRKY transcription factor during ethylene induction in Arabidopsis. PLoS Genetics, 8.

Li, H., Wang, Y., Jiang, J., Liu, G., Gao, C., & Yang, C. (2009) Identification of genes responsive to salt stress on Tamarix hispida roots. Gene, 433, 65–71.

Li, T., Chen, X., Zhong, X., Zhao, Y., Liu, X., Zhou, S., & Zhou, D. X. (2013) Jumonji C domain protein JMJ705-mediated removal of histone H3 lysine 27 trimethylation is involved in defense-related gene activation in rice. The Plant Cell, 25, 4725–4736.

Liu, Y., & Imai, R. (2018) Function of plant DExD/H-box RNA helicases associated with ribosomal RNA biogenesis. Frontiers in Plant Science, 9, 125.

Maekawa, S., Sato, T., Asada, Y., Yasuda, S., Yoshida, M., Chiba, Y., & Yamaguchi, J. (2012) The Arabidopsis ubiquitin ligases ATL31 and ATL6 control the defense response as well as the carbon/nitrogen response. Plant Mol. Biol. 79, 217–227.

Manimaran, P., Reddy, S. V., Moin, M., Reddy, M. R., Yugandhar, P., Mohanraj, S. S., et al. (2017) Activation-tagging in indica rice identifies a novel transcription factor subunit, NF-YC13 associated with salt tolerance. Scientific Reports, 7, 1–16.

Maréchal, A., & Zou, L. (2013) DNA damage sensing by the ATM and ATR kinases. Cold Spring Harbor Perspectives in Biology, 5(9), a012716.

Maselli, G. A., Slamovits, C. H., Bianchi, J. I., Vilarrasa-Blasi, J., Caño-Delgado, A. I., & Mora-García, S. (2014) Revisiting the evolutionary history and roles of protein phosphatases with Kelch-like domains in plants. Plant Physiol. 164, 1527–1541.

Mattiroli, F., Vissers, J. H., van Dijk, W. J., Ikpa, P., Citterio, E., Vermeulen, W., et al. (2012) RNF168 ubiquitinates K13-15 on H2A/H2AX to drive DNA damage signaling. Cell, 150, 1182–1195.

Maxwell, K., & Johnson, G. N. (2000) Chlorophyll fluorescence—a practical guide. Journal of Exp. Bot. 51, 659–668.

Melo, A. M., Bandeiras, T. M., & Teixeira, M. (2004) New insights into type II NAD (P) H: quinone oxidoreductases. Microbiol. Mol. Biol. Rev., 68, 603–616.

Mohanta, T. K., & Sinha, A. K. (2016) Role of calcium-dependent protein kinases during abiotic stress tolerance. Abiotic Stress Response in Plants.

Moin, M., Bakshi, A., Madhav, M. S., & Kirti, P. B. (2017) Expression profiling of ribosomal protein gene family in dehydration stress responses and characterization of transgenic rice plants overexpressing RPL23A for water-use efficiency and tolerance to drought and salt stresses. Frontiers in Chemistry, 5, 97.

Moin, M., Bakshi, A., Saha, A., Udaya Kumar, M., Reddy, A. R., Rao, K. V., Siddiq, E.A., & Kirti, P. B. (2016a) Activation tagging in indica rice identifies ribosomal proteins as potential targets for manipulation of water□use efficiency and abiotic stress tolerance in plants. Plant, Cell & Environment, 39, 2440–2459.

Moin, M., Bakshi, A., Saha, A., Dutta, M., Madhav, S. M., & Kirti, P. B. (2016b) Rice ribosomal protein large subunit genes and their spatio-temporal and stress regulation. Frontiers in Plant Science, 7, 1284.

Mok, D. W., & Mok, M. C. (2001) Cytokinin metabolism and action. Annual Review of Plant Biology, 52, 89–118.

Moon, S., & Jung, K. H. (2014) Genome-wide expression analysis of rice ABC transporter family across spatio-temporal samples and in response to abiotic stresses. Journal of Plant Physiology, 171, 1276–1288.

Mukhopadhyay, A., Vij, S., & Tyagi, A. K. (2004) Overexpression of a zinc-finger protein gene from rice confers tolerance to cold, dehydration, and salt stress in transgenic tobacco. Proceedings of the National Academy of Sciences, 101, 6309–6314.

Munns, R (1993) Physiological processes limiting plant growth in saline soils: some dogmas and hypotheses. Plant Cell Environ. 16, 15–24

Munns, R. (2005) Genes and salt tolerance: bringing them together. New Phytologist, 167, 645–663.

Munns, R., James, R. A., Xu, B., Athman, A., Conn, S. J., Jordans, C., et al. (2012) Wheat grain yield on saline soils is improved by an ancestral Na+ transporter gene. Nature biotechnology, 30, 360.

Murchie, E. H., & Lawson, T. (2013) Chlorophyll fluorescence analysis: a guide to good practice and understanding some new applications. Journal of Exp. Bot. 64, 3983–3998.

Ng, S., Giraud, E., Duncan, O., Law, S. R., Wang, Y., Xu, L., et al. (2013) Cyclin-dependent kinase E1 (CDKE1) provides a cellular switch in plants between growth and stress responses. Journal of Biological Chemistry, 288, 3449–3459.

Nishizawa, Y., Katoh, S., Koiwai, H., & Katoh, E. (2008) EL5 is involved in root development as an anti-cell death ubiquitin ligase. Plant Signaling & Behavior, 3, 148–150.

Noma, A., & Suzuki, T. (2006) Ribonucleome analysis identified enzyme genes responsible for wybutosine synthesis. In Nucleic Acids Symposium Series, 50, 65–66.

Nongpiur, R., Soni, P., Karan, R., Singla-Pareek, S. L., & Pareek, A. (2012) Histidine kinases in plants: cross talk between hormone and stress responses. Plant Signaling & Behavior, 7, 1230–1237.

Ohta, M., & Takaiwa, F. (2014) Emerging features of ER resident J-proteins in plants. Plant Signaling & Behavior, 9, e28194.

Ohta, M., Wakasa, Y., Takahashi, H., Hayashi, S., Kudo, K., & Takaiwa, F. (2013) Analysis of rice ER-resident J-proteins reveals diversity and functional differentiation of the ER-resident Hsp70 system in plants. Journal of Experimental Botany, 64, 5429–5441.

Omidbakhshfard, M. A., Omranian, N., Ahmadi, F. S., Nikoloski, Z., & Mueller-Roeber, B. (2012) Effect of salt stress on genes encoding translation-associated proteins in Arabidopsis thaliana. Plant Signaling & Behavior, 7, 1095–1102.

Pandey, R., MuÈller, A., Napoli, C. A., Selinger, D. A., Pikaard, C. S., Richards, E. J., et al. (2002) Analysis of histone acetyltransferase and histone deacetylase families of Arabidopsis thaliana suggests functional diversification of chromatin modification among multicellular eukaryotes. Nucleic Acids Res. 30, 5036–5055.

Park, J. M., Park, C. J., Lee, S. B., Ham, B. K., Shin, R., & Paek, K. H. (2001) Overexpression of the tobacco Tsi1 gene encoding an EREBP/AP2–type transcription factor enhances resistance against pathogen attack and osmotic stress in tobacco. The Plant Cell, 13, 1035–1046.

Pérez-Salamó, I., Papdi, C., Rigó, G., Zsigmond, L., Vilela, B., Lumbreras, V., et al. (2014) The heat shock factor A4A confers salt tolerance and is regulated by oxidative stress and the mitogen-activated protein kinases MPK3 and MPK6. Plant Physiol. 165, 319–334.

Petrusa, L. M., & Winicov, I. (1997) Proline status in salt-tolerant and salt-sensitive alfalfa cell lines and plants in response to NaCl. Plant Physiol. and Biochem. 35, 303–310.

Quan, R., Wang, J., Hui, J., Bai, H., Lyu, X., Zhu, Y., et al. (2018) Improvement of salt tolerance using wild rice genes. Frontiers in Plant Science, 8, 2269.

Richter, S., & Lamppa, G. K. (1998) A chloroplast processing enzyme functions as the general stromal processing peptidase. Proceedings of the National Academy of Sciences, 95, 7463–7468.

Rodnina, M. V., & Wintermeyer, W. (2009) Recent mechanistic insights into eukaryotic ribosomes. Current Opinion in Cell Biology, 21, 435–443.

Roy, S. J., Negrão, S., and Tester, M. (2014) Salt resistant crop plants. Curr. Opin. Biotechnol. 26, 115–124.

Saha, A., Das, S., Moin, M., Dutta, M., Bakshi, A., Madhav, M. S., & Kirti, P. B. (2017) Genome-wide identification and comprehensive expression profiling of Ribosomal Protein Small Subunit (RPS) genes and their comparative analysis with the Large Subunit (RPL) genes in rice. Frontiers in Plant Science, 8, 1553.

Saha, P., Mukherjee, A., & Biswas, A. K. (2015) Modulation of NaCl induced DNA damage and oxidative stress in mungbean by pretreatment with sublethal dose. Biologia Plantarum, 59, 139–146.

Sahi, C., Singh, A., Kumar, K., Blumwald, E., & Grover, A. (2006) Salt stress response in rice: genetics, molecular biology, and comparative genomics. Functional & Integrative Genomics, 6, 263–284.

Schweighofer, A., Hirt, H., & Meskiene, I. (2004) Plant PP2C phosphatases: emerging functions in stress signaling. Trends in Plant Science, 9, 236–243.

Sheen, J. (2014) Master regulators in plant glucose signaling networks. Journal of Plant Biology, 57, 67–79.

Shen, W., Wei, Y., Dauk, M., Tan, Y., Taylor, D. C., Selvaraj, G., & Zou, J. (2006) Involvement of a glycerol-3-phosphate dehydrogenase in modulating the NADH/NAD+ ratio provides evidence of a mitochondrial glycerol-3-phosphate shuttle in Arabidopsis. The Plant Cell, 18, 422–441.

Shen, Y., Conde e Silva, N., Audonnet, L., Servet, C., Wei, W., & Zhou, D. X. (2014) Overexpression of histone H3K4 demethylase gene JMJ15 enhances salt tolerance in Arabidopsis. Frontiers in Plant Science, 5, 290.

Shi, S., Li, S., Asim, M., Mao, J., Xu, D., Ullah, Z., et al. (2018) The Arabidopsis calciumdependent protein kinases (CDPKs) and their roles in plant growth regulation and abiotic stress responses. International Journal of Molecular Sciences, 19, 1900.

Shimizu-Sato, S., Tanaka, M., & Mori, H. (2009) Auxin–cytokinin interactions in the control of shoot branching. Plant molecular biology, 69, 429.

Spicher, L., Almeida, J., Gutbrod, K., Pipitone, R., Dörmann, P., Glauser, G., et al. (2017) Essential role for phytol kinase and tocopherol in tolerance to combined light and temperature stress in tomato. Journal of Experimental Botany, 68, 5845–5856.

Strizhov, N., Ábrahám, E., Ökrész, L., Blickling, S., Zilberstein, A., Schell, J., et al. (1997) Differential expression of two P5CS genes controlling proline accumulation during salt□stress requires ABA and is regulated by ABA1, ABI1 and AXR2 in Arabidopsis. Plant J. 12, 557–569.

Szklarczyk, D., Franceschini, A., Wyder, S., Forslund, K., Heller, D., Huerta-Cepas, J., et al. (2015) STRING v10: protein–protein interaction networks, integrated over the tree of life. Nucleic Acids Research, 43, 447–452.

Takatsuka, H., Umeda□Hara, C., & Umeda, M. (2015) Cyclin□dependent kinase□activating kinases CDKD; 1 and CDKD; 3 are essential for preserving mitotic activity in Arabidopsis thaliana. The Plant Journal, 82, 1004–1017.

Tamura, T., Asakura, T., Uemura, T., Ueda, T., Terauchi, K., Misaka, T., & Abe, K. (2008) Signal peptide peptidase and its homologs in Arabidopsis thaliana–plant tissue□specific expression and distinct subcellular localization. The FEBS Journal, 275, 34–43.

Tester, M., & Davenport, R. (2003) Na+ tolerance and Na+ transport in higher plants. Annals of Bot. 91, 503–527.

Tohge, T., Watanabe, M., Hoefgen, R., & Fernie, A. R. (2013) Shikimate and phenylalanine biosynthesis in the green lineage. Frontiers in Plant Science, 4, 62.

Tsukada, Y. I., Fang, J., Erdjument-Bromage, H., Warren, M. E., Borchers, C. H., Tempst, P., & Zhang, Y. (2006) Histone demethylation by a family of JmjC domain-containing proteins. Nature, 439, 811–816.

Verma, S., & Mishra, S. N. (2005) Putrescine alleviation of growth in salt stressed Brassica juncea by inducing antioxidative defense system. Journal of plant physiol. 162, 669–677.

Voss, M., Schröder, B., & Fluhrer, R. (2013) Mechanism, specificity, and physiology of signal peptide peptidase (SPP) and SPP-like proteases. Biochimica Et Biophysica Acta (BBA)-Biomembranes, 1828, 2828–2839.

Wang, K., Wang, M., Tang, D., Shen, Y., Miao, C., Hu, Q., et al. (2012) The role of rice HEI10 in the formation of meiotic crossovers. PLoS Genetics, 8(7).

Wu, H. (2018) Plant salt tolerance and Na+ sensing and transport. The Crop Journal, 6, 215–225.

Xiang, Y., Huang, Y., & Xiong, L. (2007) Characterization of stress-responsive CIPK genes in rice for stress tolerance improvement. Plant Physiol. 144, 1416–1428.

Xing, Z., Ma, W. K., & Tran, E. J. (2019) The DDX5/Dbp2 subfamily of DEAD□box RNA helicases. Wiley Interdisciplinary Reviews: RNA, 10, e1519.

Xue, T., Wang, D., Zhang, S., Ehlting, J., Ni, F., Jakab, S., et al. (2008) Genome-wide and expression analysis of protein phosphatase 2C in rice and Arabidopsis. BMC Genomics, 9, 550.

Yang, C., Zang, W., Ji, Y., Li, T., Yang, Y., & Zheng, X. (2019) Ribosomal protein L6 (RPL6) is recruited to DNA damage sites in a poly (ADP-ribose) polymerase–dependent manner and regulates the DNA damage response. Journal of Biological Chemistry, 294, 2827–2838.

Yasuda, S., Sato, T., Maekawa, S., Aoyama, S., Fukao, Y., & Yamaguchi, J. (2014) Phosphorylation of Arabidopsis ubiquitin ligase ATL31 is critical for plant carbon/nitrogen nutrient balance response and controls the stability of 14-3-3 proteins. Journal of Biological Chemistry, 289, 15179–15193.

Zhang, Z., Zhang, Q., Wu, J., Zheng, X., Zheng, S., Sun, X., et al. (2013). Gene knockout study reveals that cytosolic ascorbate peroxidase 2 (OsAPX2) plays a critical role in growth and reproduction in rice under drought, salt and cold stresses. PloS One, 8(2).

Zhao, L., Li, Y., Xie, Q., & Wu, Y. (2017) Loss of CDKC; 2 increases both cell division and drought tolerance in Arabidopsis thaliana. The Plant Journal, 91, 816–828.

Zheng, T., Nibau, C., Phillips, D. W., Jenkins, G., Armstrong, S. J., & Doonan, J. H. (2014) CDKG1 protein kinase is essential for synapsis and male meiosis at high ambient temperature in Arabidopsis thaliana. Proceedings of the National Academy of Sciences, 111, 2182–2187.

